# *In vitro* cell cycle oscillations exhibit a robust and hysteretic response to changes in cytoplasmic density

**DOI:** 10.1101/2021.05.19.444890

**Authors:** Minjun Jin, Franco Tavella, Shiyuan Wang, Qiong Yang

**Author notes:** These authors contributed equally to this work (Minjun Jin, Franco Tavella).

## Abstract

Cells control the properties of the cytoplasm to ensure the proper functioning of biochemical processes. Recent studies showed that the density of the cytoplasm varies in both physiological and pathological states of cells undergoing growth, division, differentiation, apoptosis, senescence, and metabolic starvation. Little is known about how cellular processes cope with these cytoplasmic variations. Here we study how a cell cycle oscillator comprising cyclin -dependent kinase (CDK1) responds to cytoplasmic density changes by systematically diluting or concentrating a cycling *Xenopus* egg cytoplasm in cell-like microfluidic droplets. We found that the cell cycle maintains robust oscillations over a wide range of deviations from the endogenous density by as low as 0.2x to more than 1.22x. A further dilution or concentration from these values will arrest the system in a low or high steady-state of CDK1 activity, respectively. Interestingly, diluting a concentrated arrested cytoplasm recovers its oscillatory behavior but requires a significantly lower concentration than 1.22x. Thus, the cell cycle switches reversibly between oscillatory and stable steady states at distinct thresholds depending on the direction of density tuning, forming a hysteresis loop. We recapitulated these observations by a mathematical model. The model predicted that Wee1 and Cdc25 positive feedback do not contribute to the observed robustness, confirmed by experiments. Nevertheless, modulating these feedback strengths and cytoplasmic density changes the total number of cycles, revealing a new role of Wee1 and Cdc25 in controlling the cycle number of early embryonic extracts. Our system can be applied to study how cytoplasmic density affects other cellular processes.

## INTRODUCTION

Robustness, the ability to maintain function despite environmental perturbations, appears ubiquitous across many areas of biology. Processes as diverse as bacterial chemotaxis, osmoregulation, circadian rhythms, and development have been found insensitive to certain conditional changes in a wide range (1-4). Rather than requiring precise tuning of parameters, studies have attributed the stability of their functioning to a robust property of their underlying biochemical networks.

At the cellular level, biochemical reactions take place in a highly complex and dynamic environment called the cytoplasm. Activities including synthesis, degradation, nucleocytoplasmic translocation, osmosis, endocytosis, and exocytosis may all impose perturbations to the state of a eukaryotic cytoplasm. It is crucial to understand how the cytoplasmic properties vary and how the internal processes respond to these fluctuations and remain stable functioning.

Recent advances in precise cell density measurement, particularly by the suspended microchannel resonator (SMR) and quantitative phase microscopy, have revealed significant changes in the overall cytoplasmic density throughout the cell cycle (5-8). Different mammalian cell types display a common phenomenon of mitotic swelling with a significant volume increase (10-30%) and a constant dry mass, resulting in a drop of cell density during mitosis (7,8).

Additionally, it has been shown that large deviations from the physiological ranges of cytoplasmic density can induce detrimental cell states such as senescence and apoptosis (9-13). Interestingly, some processes remain unaffected despite considerable variations in the density of the cytoplasm. For example, after a 25% loss of intracellular water, the glucose metabolism in rat liver cells shows no noticeable change in its reaction rates (14). Thus, cells must tolerate some degree of cytoplasmic density changes but can respond to severe deviations from the physiological state. Although the cytoplasmic density has gained increasing recognition as an essential parameter to control proper cellular function, it remains unclear how fundamental cellular processes, such as the cell cycle, behave under variations in cytoplasmic density.

Computational studies have demonstrated the capability of cell-cycle networks to perform robust functions. The yeast cell-cycle network shows persistent dynamical properties against small perturbations to the network topology (15). Another study associated the positive-plus-negative feedback architecture at the core of the embryonic cell-cycle network with the increased robustness to changes in the system’s parameters (16). But to date, there has been no computational or experimental work that shows how the cell-cycle network would react to the dilution or concentration of its components as the cytoplasm fluctuates. It is also experimentally challenging to flexibly manipulate the cytoplasmic density in live cells without sabotaging other essential functions.

Here we address this question by analyzing how *in vitro* cell cycle dynamics change with the density of *Xenopus* cell-free cytoplasmic extracts in a microfluidic droplet system. We used programmed pressure-driven control of inlet flow in a microfluidic device to generate droplets encapsulating extracts with different dilutions. We then measured the period and persistence of oscillations in these droplets using a CDK1 activity FRET sensor. We found that cell cycle oscillations withstand both dilution and concentration for an extensive range of cytoplasmic densities. We further developed a mathematical model to investigate the role of cytoplasmic density in the oscillatory behavior of the cell-cycle network. Finally, we explored whether Wee1 and Cdc25, the key components underlying the positive-plus-negative feedback architecture, are necessary for the robustness of the cell cycle to cytoplasmic density changes.

## RESULTS

### *In vitro* cell cycle oscillations are robust to cytoplasmic dilutions

We first reconstituted mitotic oscillation *in vitro* using cycling *Xenopus* cytoplasmic extracts. The system periodically alternates between interphase and mitotic phases by a self-sustained cell-cycle oscillator centered on the cyclin-dependent kinase (CDK1). The phosphorylation– dephosphorylation cycle of mitotic substrates is co-regulated by the protein kinase CDK1-cyclin B and its antagonistic phosphatase PP2A-B55δ, which mutually regulate each other’s auto-activation loops (Fig 1A). To measure the activity ratio between CDK1-cyclin B and PP2A-B55δ in extracts, we applied a recently developed CDK1 FRET sensor (G.M. and Q.Y., unpublished work), which contains a mitotic substrate Cdc25C-ligand domain and phosphorylation-sensing WW domain. When extracts are in a high CDK1-cyclin B activity state, phosphorylated groups on the Cdc25c-ligand domain are detected by sensing the WW domain, bringing the donor, cyan fluorescent protein (CFP), and the acceptor, yellow fluorescent protein (YFP), in proximity for energy transfer.

**Figure 1.**
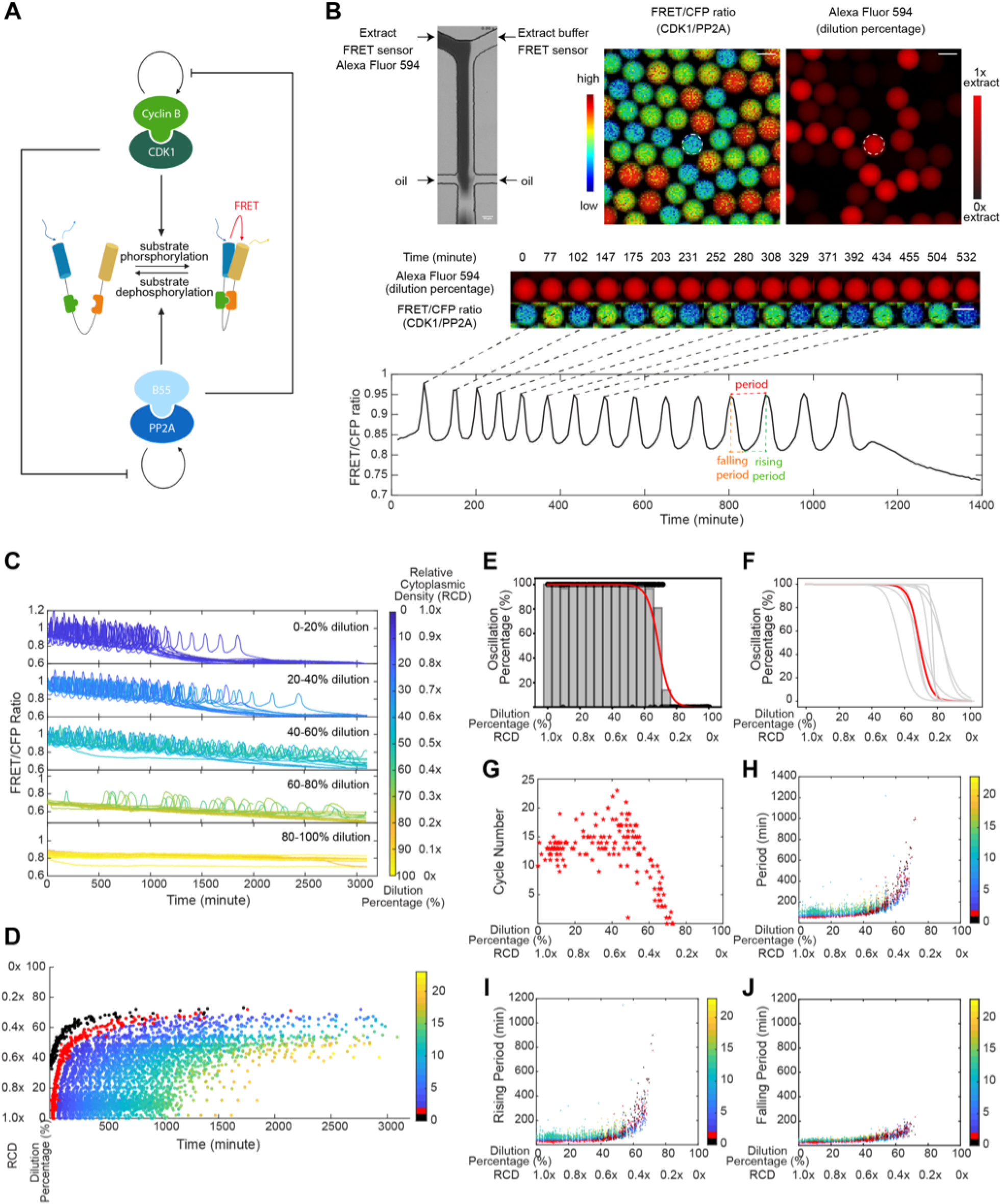
In vitro cell-cycle oscillators are robust to cytoplasmic dilutions. **(A)** Schematic diagram of the phosphorylation and de-phosphorylation cycles of mitotic substrates driven by kinase, CDK1-cyclin B and its counteracting phosphatase, PP2A-B55, which mutually modulate each other’s auto-activation loops. The full diagram for the cell-cycle oscillator is in Fig 3A. A CDK1 FRET sensor is used to measure the ratio of CDK1-cyclin B to PP2A:B55 activity throughout oscillations. The sensor is composed of a Cdc25C-ligand domain and a phosphorylation-sensing WW domain. When the ligand domain is phosphorylated, the affinity for the sensing domain increases, bringing the donor and acceptor fluorescent domains together and resulting in increased FRET efficiency. **(B)** FRET-based quantification of cell cycle dynamics in microemulsion droplets containing 0-100% diluted *Xenopus* extracts. **Top left:** A microfluidic device generates droplets containing extracts and Extract buffer mixed at different ratios. The extracts supplied with Alexa Fluor 594 Dextran dye flow through one inlet and the extract buffer through the other, both containing the same concentration of CDK1-FRET sensor. The diluted extracts are then encapsulated by surfactant oil into uniform-sized micro-emulsion droplets. **Top right:** Snapshots of FRET/CFP ratio and Alexa Fluor 594 Dextran channels for a field of droplets indicating the ratio of CDK1/PP2A activity and the dilution percentage. **Middle:** Time-lapse series of a selected droplet showing the first 550-minute imaging in both channels. Selected time points correspond to either a peak or trough in FRET/CFP signals. The droplet has a constant red fluorescence intensity over time, indicating no leakage of the dye. The scale bar is 100 μm. **Bottom:** Time course of FRET/CFP ratio of the selected droplet showing 15 undamped oscillations in 1400 minutes. The oscillation period is quantified as the time interval between two consecutive peaks. The rising period (interphase duration) is defined as the time interval between a trough to the subsequent peak, and the falling period (duration of mitosis) as the time interval between a peak to the subsequent trough. **(C)** Time courses of FRET/CFP ratio for droplets with 0-100% diluted extracts show the cell cycle robustness to cytoplasmic dilutions. Each line represents a droplet, and the color of the line indicates the dilution percentage (relative cytoplasmic density, RCD) of the droplet. The cytoplasmic density of undiluted droplets (dilution 0%) is defined as 1.0x. Twenty droplets are randomly selected from each dilution range and presented in the plot. **(D)** Raster plot of the onset of FRET/CFP ratio peaks over time shows the entry of the first cycle is significantly delayed as dilution percentage increases, and the overall oscillation time (from the start of the first peak to the last peak) is extended as well. Each dot represents a cycle peak in a droplet. The color bar indicates the order of peaks. Here we define the first observed cycle of undiluted extracts as cycle 1 (labeled in red). When extracts have been more diluted, we observe the delayed cycle 0 (labeled in black). **(E)** Oscillation percentage of droplets over different dilution ranges. Oscillation percentage is calculated as the percentage of droplets that exhibit oscillations in all tracked droplets within each bin range. The bin width in the histogram is 5. The red curve is the logistic regression fitting result for oscillation percentage data. This dataset includes 655 detected droplets, in which 351 droplets yield oscillations. **(F)** Logistic regression fits for oscillation percentage versus dilution for nine replicates, showing the cell cycle robustness to dilutions is highly conserved across all different day experiments. The red highlighted curve corresponds to the experiment in panel E. **(G)** The cycle number versus dilution of individual droplets presents an initial plateau within low-moderate dilution ranges, and subsequently decreases as a function of dilution. Cycle number for each droplet was quantified by a total number of peak-to-peak FRET/CFP oscillations. **(H)** Oscillation periods remain relatively constant up to 50% dilution and afterward increase dramatically with dilution until the dilution is close to 80%, when oscillation arrests at a low CDK1 activity. Similar trends are observed in **(I)** rising periods and **(J)** falling periods. Each dot indicates one cycle in a droplet. The color bar represents the order of cycles.

To study the effect of cytoplasmic dilution on cell cycles, we encapsulated the cytoplasmic extracts into water-in-oil microemulsions (17,18) and employed microfluidic devices with two inlets to perform dilutions. Programmed pressure-driven control of the flow in the inlets continuously adjusts ratios of cytoplasmic extract and extract buffer in individual droplets, resulting in a full spectrum of cytoplasmic dilution from 0 to 100% (1.0x to 0x relative cytoplasmic density, RCD) (Fig 1B, Top). Each droplet was supplied with 1 μM CDK1 FRET sensor to track the oscillation progression and a fluorescent dye (Alexa Fluor 594) that co-injects with the extracts into one inlet to quantify the dilution percentage. These droplets were loaded into Teflon-coated glass tubes, recorded for up to 3 days using time-lapse fluorescence microscopy, and analyzed for their FRET/CFP ratio time courses to extract oscillation parameters (Fig 1B, Middle and Bottom; Movie S1).

Surprisingly, droplets retain oscillations for dilutions as large as 80% (0.2x RCD) (Fig 1C). The overall waveform was invariant up to 40% dilutions (0.6x RCD), with the remaining range from 40%-80% showing longer rising and falling phases. Nevertheless, the system settled into a stable steady-state of CDK1 when dilution was in the range of 80%-100%. We recapitulate these trends in a raster time series of oscillation peaks of all individual droplets sorted by their dilution percentage (Fig 1D). It further reveals that the start and end of oscillations in each droplet are also dilution dependent. As the dilution percentage increases, both the entry of the first peak and the peak time of the last cycle were delayed, resulting in an overall longer time window of oscillation. The phenomenon was more pronounced when extracts were greatly diluted (above 40%, 0.6x RCD).

To quantify the impact of cytoplasmic dilution on mitotic oscillations, we calculated the percentage of oscillatory droplets within each dilution range (Fig 1E). Between 0% and 60%, all tracked droplets showed oscillations. Close to 80%, the oscillation percentage dropped abruptly to zero. This observation was well conserved across nine independent experiments (each performed on a different day), except for slightly different absolute half-maximal thresholds (defined by the dilution percentage at which 50% of the droplets oscillate), which are likely due to batch variations in *Xenopus* egg extracts (Fig 1F). These results demonstrated the oscillation robustness to cytoplasmic dilution.

Additionally, we observed that the total number of cycles in each droplet presented an initial plateau, followed by an abrupt decrease near 50% dilution (0.5x RCD) (Fig 1G). We also analyzed three other properties of the oscillations, namely the total period (Fig 1H), rising period (duration of interphase) (Fig 1I), and falling period (duration of M phase) (Fig 1J). The total period of oscillations remained relatively unchanged until 50% dilution when it increased abruptly. Rising and falling periods followed a similar trend as the total period, although the rising periods showed a greater fold change at a given dilution than the falling ones (Fig S1). This differential sensitivity of rising and falling phases to variations has been previously reported in other studies, where mitosis seems temporally insulated from variability in other cell-cycle events (19,20). In addition, we observed batch variations in the period and cycle-number responses to dilution across different day experiments (Fig S2), emphasizing the influence of the initial egg conditions on the robustness of mitotic oscillators. Nevertheless, the overall trend in the period was conserved for all replicates. These results indicated that the system not only is robust in its ability to oscillate but maintains invariant oscillation profiles for a broad range of dilutions.

To test whether the behavior near the transition threshold could be solely explained by partitioning variability during encapsulation, we generated droplets with two different diameters of 83.18±2.21 μm and 55.29±1.83 μm (Fig S3A). If the halt of oscillations was caused by the uneven distribution of cytoplasmic material, smaller droplets would be more impacted by partition errors and display a lower oscillation threshold and a larger variation in the high dilution regions. However, we did not observe a significant difference between the two cases when we compared their oscillation percentage curves (Fig S3B), distribution of period lengths (Fig S3C), and total cycle numbers (Fig S3D). Thus, the behavior of the system near the transition threshold was inherent to the response of the mitotic circuit to changes in cytoplasmic density rather than caused by an uneven distribution of cytoplasmic material.

### Cytoplasmic concentration leads to a reversible loss of oscillation and displays hysteresis

Next, we investigated the effect of concentrating the cytoplasm by vacuum evaporation of the bulk cycling *Xenopus* extract before encapsulation. We used the fluorescent intensity changes of Alexa Fluor 594 to quantify the resultant relative cytoplasmic density after concentration, confirmed by other measurements using Nanodrop and volume (Table S1). We found the system sustained oscillations with no noticeable change for extracts with evaporation time up to 30 minutes (1.22x RCD) (Fig 2A and 2B, Row 1-4; Movie S2). When further concentrating the extract for 40 minutes to 1.46x RCD, the system showed no oscillations (Fig 2A and 2B, Row 5). The results suggested a threshold of RCD between 1.22x and 1.46x, beyond which cell cycle arrests, in this case, at a high stable state of CDK1 compared to the stably low CDK1 activity for a highly diluted system (Fig 1C, Row 5). These two distinct CDK1 steady states implied there exist different mechanisms to regulate the halt of oscillations at the two cytoplasmic density extremes.

**Figure 2.**
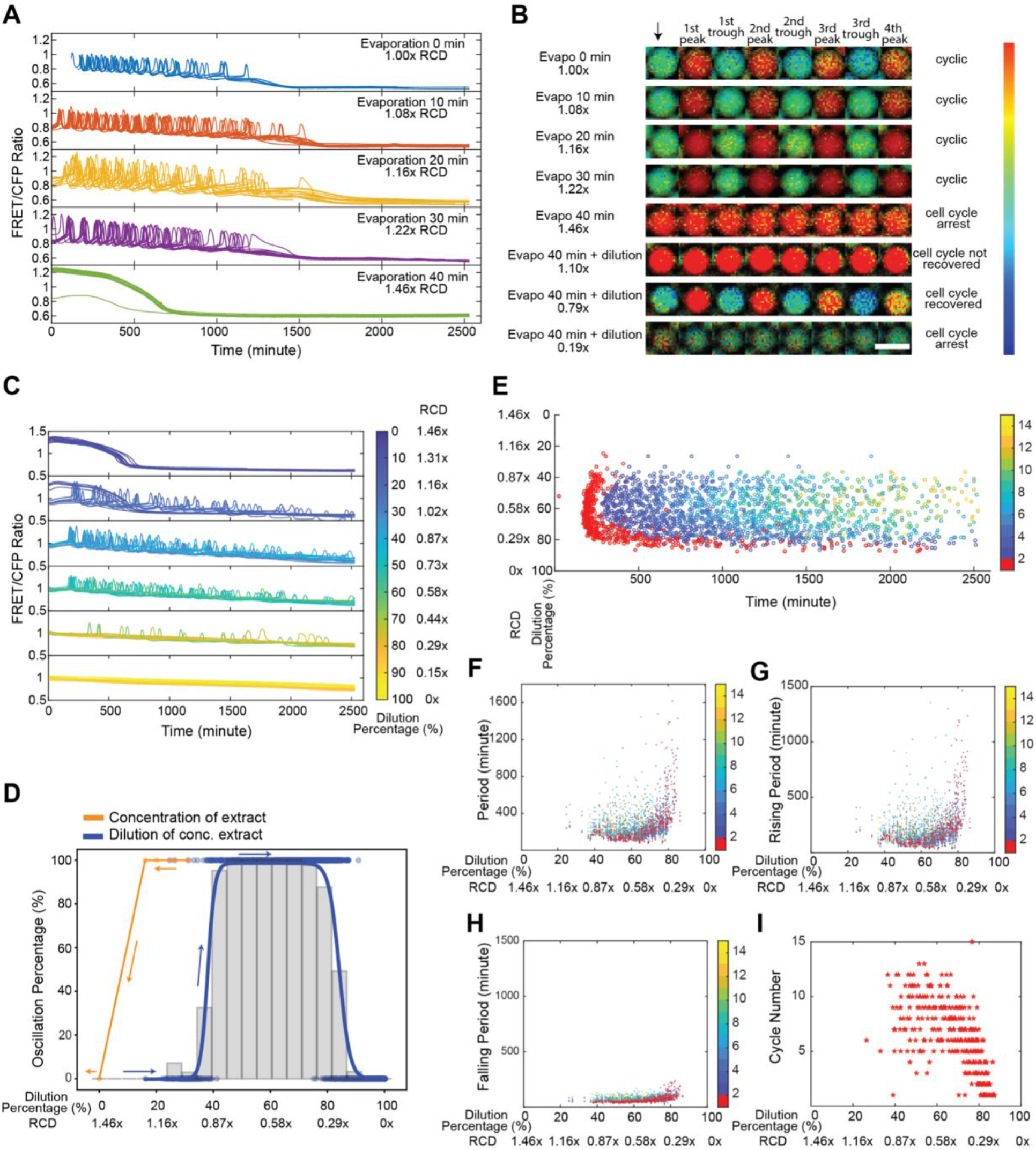
Cytoplasmic concentration leads to a reversible loss of oscillation but displays hysteresis. **(A)** Time courses of FRET/CFP ratio for individual droplets with concentrated extracts, showing cell cycles are robust to cytoplasmic concentrations. Extracts are concentrated via vacuum evaporation with the resultant relative cytoplasmic densities quantified by the intensity changes in Alexa Fluor 594 dextran dye before and after the concentration process. We define the cytoplasmic density of undiluted extracts as 1x, and the RCD for extracts with 10, 20, 30, and 40-min evaporation are measured to be 1.08x, 1.16x, 1.22x, and 1.46x, respectively. For ease of visualization, twenty droplets are randomly selected from each condition and presented in the plot. The sample size for different conditions (from top to bottom) is 46, 471, 140, 98, and 506, respectively. **(B)** FRET/CFP ratio images of representative droplets with varying cytoplasmic densities by initially concentration (from the 1^st^ row to the 5^th^ row) then dilution (from 5^th^ row to 8^th^ row), which demonstrates dilution of non-oscillatory concentrated extracts restores oscillations. The arrow indicates the start of imaging. Each following image is associated with either a trough or a peak of the selected droplets. Droplets with 1.00x, 1.08x, 1.16x, and 1.22x RCD present normal oscillations of CDK1. The droplet with 40-min evaporation (1.46x RCD) stays at a high CDK1 activity without any noticeable periodic behavior, suggesting the system has settled to a steady state. The addition of Extract buffer into the non-oscillatory 1.46x extracts to achieve 1.10x RCD cannot recover oscillations, but oscillations only reappear in the droplet with a dilution down to 0.79x RCD. Further dilution of the system to as low as 0.19x eventually arrests the cell cycle again, this time at a low CDK1 activity steady-state, as compared to the high steady-state of CDK1 in the 1.46x concentrated extracts. The scale bar is 100 μm. The data for dilutions of concentrated extracts contains two replicates, resulting in a sample size of 1198 detected droplets and in which, 538 droplets have at least one cycle. **(C)** Time courses of FRET/CFP fluorescence intensity ratio in droplets containing 1.46x concentrated extracts with different dilutions, showing clear transitions from a high CDK1 activity arrested state to oscillations to a low CDK1 state. As the 1.46x concentrated extracts are being diluted, droplets with low dilutions (0-30%) tend to stay at a high steady state of CDK1. Droplets with higher dilutions (30%-60%) restore oscillations. Further dilutions (>70%-80%) lead to a cell cycle arrest at a low steady state of CDK1. Each line is for one droplet, and the color of the line indicates the dilution percentage of the droplet. Twenty droplets are shown for each dilution range. Here, zero dilution has 1.46x RCD. **(D)** Cell cycle shows a hysteretic response to the change of cytoplasmic density. The change in oscillation percentage follows the orange curve when we concentrate the cytoplasmic density from 1.00x to 1.46x. The system transitions from limit cycles to a stable steady-state, at a threshold value in between 1.22x and 1.46x. When we start to dilute the concentrated extracts at 1.46x RCD, the oscillation percentage changes along the blue curve. These two distinct paths show a “history dependent” nature. **(E)** Raster plot for the dilution of concentrated extract experiment. The first peak onset has larger variations for droplets lying at the highly diluted threshold end (∼80%). The color bar indicates the order of cycle peaks, and the peak of the first cycle is highlighted in red. **(F)** Periods remain relatively constant at moderate dilutions after restoring oscillations (following the blue curve in Fig 2E), with the RCD approximately 0.80x to 0.44x. The variations are significant near two ends, especially the diluted end (0.29x RCD). Similar trends are observed in **(G)** rising periods and **(H)** falling periods. Each dot represents a cycle in a droplet. The color bar indicates the order of cycles, and the first cycle is highlighted in red. **(I)** Droplets with moderate dilutions have more cycles than those with either too low or too high dilution percentages at two ends.

Nevertheless, as we diluted the 1.46x concentrated extract with 2-inlet microfluidics, oscillation reappeared, and we observed a clear transition from a stable high CDK1 steady-state to limit cycle oscillations to a stable low CDK1 steady-state (Fig 2B, Row 6-8; Fig 2C; Movie S3). Thus, the oscillatory behavior can be modulated in a reversible manner by cytoplasmic density. Additionally, by diluting the 1.46x concentrated extract, the oscillation percentage of droplets recovered from 0 to 100% around 40% dilution (0.87x RCD) and then back to 0 at a high dilution of 90% (Fig 2D, blue curve), defining the two boundaries of the distribution. Compared to the threshold to stop oscillations (in between 1.22x and 1.46x RCD) when concentrating the extracts (Fig 2D, orange curve), the same system cannot restore oscillations by diluting until below a threshold of about 0.87x RCD. Therefore, the transitions between the two states, i.e., oscillatory state and a high CDK1 stable steady-state, do not follow the same path as the system undergoes concentration versus dilution, suggesting the possibility of hysteresis.

Over the entire dilution range, the system displayed high variations in the onset of peaks near the two oscillation boundaries of 40% and 90% dilutions (Fig 2E). The highest variations were near 90% dilution, comparable to the near-boundary behavior of the endogenous-extract dilution (Fig 1D). For droplets with intermediate RCD ranging from 0.80x to 0.44x, peaks occurred at relatively similar times. We observed a similar trend for the total period (Fig 2F), rising period (Fig 2G), and falling period (Fig 2H). The total number of cycles reached a maximum when the RCD was approximately 0.80x to 0.44x and decreased at both ends of cytoplasmic densities (Fig 2I), suggesting the less tendency of the system to oscillate near two boundaries, as also observed in oscillation percentage distribution (Fig 2D, blue curve).

### Mathematical modeling recapitulates the robustness to changes in cytoplasm density and its hysteretic response

Modifying cytoplasmic density alters the kinetics of a system at the molecular level (5). To further explore the mechanisms behind the observed robustness, we developed a mathematical description of the effect of cytoplasmic density on cell cycle dynamics. We started from a previously published model of *Xenopus* early embryonic cycles that describes the dynamics of the network’s two master regulators (CDK1-cyclin B1 and PP2A-B55δ) (21). Fig 3A (Top panel) shows a diagram of the principal biochemical reactions and the influence between proteins. Briefly, cell cycles are driven by the synthesis and regulated degradation of cyclinB1. Additional layers of control provide bistability to the activity of the master regulators: kinase CDK1-cyclin B1 complexes switch between a high and low state thanks to the positive feedback regulation of Cdc25C and Wee1A; similarly, the phosphatase PP2A-B55δ alternates between high and low activity thanks to the positive feedback regulation created by Greatwall kinase and the stoichiometric inhibitor ENSA. The pathways of both master regulators are interconnected through mutual inhibition of each other’s positive feedback. When CDK1-cyclin B1 reaches its high activity state, proteolytic degradation of cyclinB1 is activated in an ultrasensitive way through the activated APC/C by CDK1. Cyclin B1 degradation restarts the system for a new cycle.

**Figure 3.**
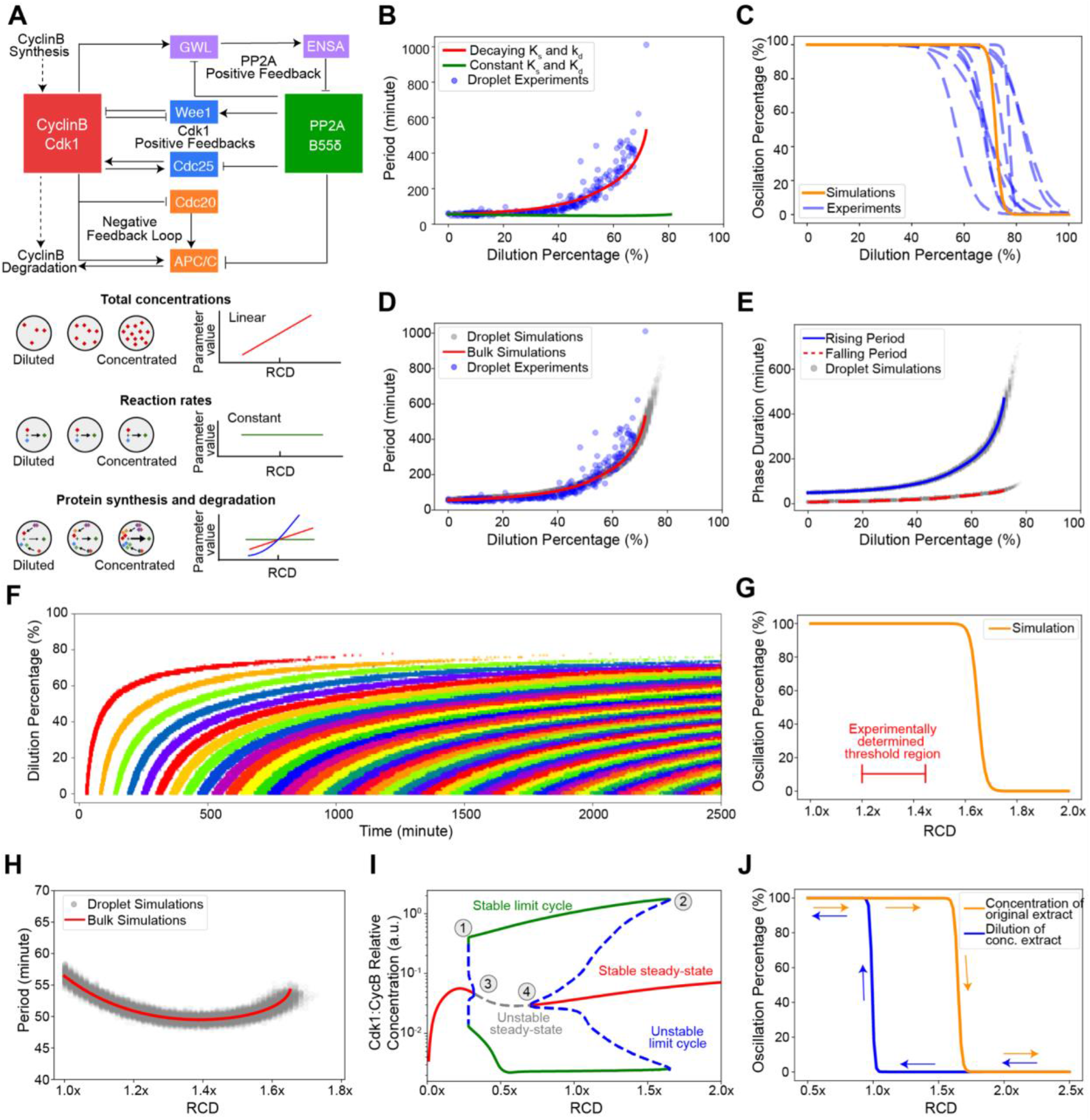
Dynamical model of the cell cycle reproduces the observed robustness of oscillations and the hysteretic response to cytoplasmic density. **(A)** Schematic view of the network controlling cell cycle progression and dependency of parameters with cytoplasmic density. **Top:** Each solid arrow corresponds to an influence present in the model. Pointed-headed arrows indicate activation and blunt-headed arrows inhibitions. Activation/Inhibitions arise from phosphorylation/dephosphorylation reactions or binding/unbinding events. Dashed arrows represent synthesis and degradation processes. **Bottom:** Different dependencies with cytoplasmic density are assumed for the parameters of the model. Total concentrations of proteins are considered to scale linearly with cytoplasmic density. In contrast, phosphorylation/dephosphorylation and binding/unbinding rates are considered to be constant because they are considered elementary reactions. Dependency of synthesis and degradation rate on cytoplasmic density are more complex as many reactions underlie their effective reaction rate, and are thus determined by comparing model predictions with experimental observations. **(B)** Period of oscillations as a function of dilution percentage. Comparison between models with decaying or constant synthesis/degradation rates. The decaying model has ks ∼ d^3^ and kd ∼ d^2^ with d the RCD. Both models include a linear scaling of total concentrations with dilution. The model without decaying rates does not feature an increase in periods consistent with experimental observations. **(C)** A sigmoidal fit to the simulated oscillation percentage of droplets versus dilution is shown as a solid orange curve. Experimental curves (dashed blue line) are repeated from Fig 1F for comparison. At a threshold of close to 80%, most droplets settle into a steady-state consistent with experimental observations. **(D)** Simulated period as a function of dilution for droplet (gray dots) and bulk (red line) models. Bulk simulations are performed with a single set of ODE parameters representing the droplet’s average parameters. Experimental data (blue dots) is presented for comparison. Both simulated and experimentally determined periods follow a similar trend with a slow increase at first and a sudden increase close to the threshold. **(E)** Duration of the rising and falling phases of the oscillation. Solid blue and dashed red lines represent bulk simulations for rising and falling periods respectively. Dots represent droplet simulations. For both types of simulations, the rising period is more affected than the falling period. **(F)** Raster plot of the oscillation peaks over time for different dilution values. Each cycle is colored differently for visual clarity. Simulations show a similar trend for the start of oscillations observed in experiments. **(G)** Oscillation percentage from droplet simulations when the RCD is increased. The experimentally determined threshold is smaller than the one obtained through simulations. **(H)** Period as a function of RCD for concentration of extracts. Bulk and droplet simulations are shown. For the concentration of the extract, the period remains comparatively constant with respect to dilution. **(I)** Bifurcation diagram with RCD as a control parameter. Vertical axis shows the maximum and minimum of Active CDK1:CyclinB1 concentration. Stable dynamical states are represented by solid lines and unstable ones by dashed lines. Each color represents a different dynamical behavior. The stable limit cycle solution disappears through a saddle-node bifurcation by meeting with an unstable limit cycle solution (1 and 2). Two supercritical Hopf-bifurcations (3 and 4) are observed for both concentration and dilution of the extract. This bifurcation structure leads to a hysteretic behavior in the appearance/disappearance of oscillations. **(J)** Hysteresis in the onset of limit cycle oscillations. Two different initial conditions are used for the simulations of each curve. One that is closer to the limit cycle and another one closer to the steady-state in the concentrated extract. All time series are obtained by numerical integration of the ODE equations. Oscillatory or steady-state behavior is determined by the norm of the system’s velocity-vector in conjunction with an analysis of the signal’s peak detection features. Namely, the period stability (variability in time differences between detected peaks) and amplitude stability (variability in peak concentrations) are analyzed. Droplet simulations are ODE simulations where the corresponding total concentrations, synthesis rate, and degradation rate are sampled from a Gamma distribution representing the encapsulation error. In all cases, 500 droplets are simulated per relative cytoplasmic density value.

Dilution and concentration can change the underlying parameters that govern the biochemical reactions of a dynamic system (Fig 3A, Bottom panel). Here, we introduced a single scaling factor in our model to characterize the changes in cytoplasmic density and scale all parameters appropriately. In particular, we first assumed the macromolecular concentrations scale linearly with the cytoplasmic density. Second, we assumed that the rates of fundamental reactions such as phosphorylation/dephosphorylation and binding/unbinding events are not directly affected by changes in cytoplasmic density. Nevertheless, it is important to note that phosphorylation/dephosphorylation and binding/unbinding reactions will be indirectly affected by changes in the total concentration of the reactants. This indirect effect was taken care of via the scaling of total concentrations but not directly through modifications of the reaction rates. Lastly, two major processes, i.e., synthesis and degradation of cyclin B1, involve multiple proteins and complex interactions and are very likely to be affected by changes in cytoplasmic density. However, the detailed reactions, such as ribosome-mRNA interactions and ubiquitin-proteasome pathways, are not explicitly modeled in our present level of description. To obtain an appropriate scaling function for both the synthesis and degradation rates, we simulated the change in the period as a function of cytoplasmic dilution with different polynomial scalings of both rates (Fig S4, SI text S2). While we could not recapitulate experimental observations in our model if the synthesis and degradation rates do not vary with density, we found the best match of simulation results with experimentally observed trends when both synthesis and degradation decay with dilution: oscillations remain up to 80% and the period increases abruptly after 60% (Fig 3B, Fig S5, SI text S2). Therefore, our results suggest that the main driving force for the observed behavior is the cytoplasmic-density-dependent changes in synthesis and degradation rates, and the decay of total concentrations alone could not explain the observed large increase in period (Fig 3B, green curve).

Biochemical reactions encapsulated in cell-sized volumes can show different behaviors than their bulk counterparts (22). Not only the absolute number of molecules is different in each case, but the process of droplet encapsulation usually implies a compositional variability between compartments. To explain the experimentally observed variability in the period and likelihood of oscillation among droplets with the same cytoplasmic density, we considered the partitioning effects in our model and simulated a population of *in silico* droplets. Each droplet draws its parameters from a gamma distribution which represents the partition errors that occur when the cytoplasm is encapsulated (SI text S3, Fig S6). This broader-than-Poisson distribution has been previously shown to accurately capture the multiple factors affecting a compartment’s composition in the process of encapsulation (22). Additionally, we used the average of each parameter in a deterministic model to describe the bulk reaction (Fig S6, red curves).

After simulating a population of droplets for each dilution, we observed that the ensemble probability of oscillation follows the same trend as experiments with a threshold within the range of the experimentally measured values (Fig 3C). We also observed an agreement of the period to dilution dependency among *in silico* droplets, bulk simulation, and experiments (Fig 3D). Interestingly, droplet simulations extend beyond the threshold set by the bulk system, highlighting the importance of considering single-cell variability in sensitive zones of parameter space. Rising and falling phases of the oscillation also show the same trend as experiments with the rising periods more sensitive to large dilutions than the falling ones (Fig 3E). Notably, droplet simulations also showed the same trends in the dispersion of the period as experiments. Namely, the period is less dispersed for low dilutions (less than 5 minutes) than near the threshold, when the period varies within a range of 45 minutes. Finally, when we simulated our system with the same initial condition of low CDK1-cyclin B1 activity for all dilutions, we obtained a delay in the onset of oscillations similar to the trend observed in experiments (Fig 3F). This delay and the subsequent period lengthening were likely caused by low cyclin B1 synthesis rates at high dilution percentages. In conclusion, the agreement between experimental features and simulations suggests that our current understanding of the cell cycle network and the assumed effect of dilution on the model’s parameters can account for the observed robustness of the system.

Next, we explored how concentrating the components of a system affects its properties. For parsimony, we assumed that the same scaling functions that ruled the effect of dilution also explain the behavior during density increase. We observed that a population of simulated droplets stops oscillating as the cytoplasmic density passes a threshold value (Fig 3G). The predicted threshold is overestimated compared to the experimentally measured values, which may be because both synthesis and degradation rates become saturated after certain concentrations rather than continue to increase polynomially as in our proposed scaling. Nevertheless, the period of these oscillations does not present large deviations, consistent with experiments, underscoring the robustness of the oscillator (Fig 3H). We summarized the complete dynamical behavior of the system in a bifurcation diagram (Fig 3I). As cytoplasmic density increases or decreases, stable oscillatory solutions (green curve) eventually disappear (points 1 and 2) and the system settles into a steady-state (red curve). Intriguingly, the bifurcation diagram presents a much richer dynamical phenomenon: the coexistence of multiple stable solutions for a specific cytoplasmic density. For example, when cytoplasmic density is 1.25x, the system can either be oscillating or in a stable steady-state, depending on the initial condition. This is evidenced by the subcritical Hopf bifurcations present in the diagram (points 3 and 4). In addition, the existence of this multistability provides a hysteretic transition between oscillatory and steady-state behaviors. If the system starts from an initial condition that promotes oscillations, as the cytoplasmic density increases, oscillations stop close to 1.7x. However, if the system starts from an initial condition that promotes a stable steady-state, the threshold is closer to 1.0x. This is shown in simulations where the hysteresis can be explicitly observed (Fig 3J). These results suggest an explanation for the different thresholds observed experimentally when concentrating and diluting back the system.

### Cdk1 positive feedbacks are not essential for robustness to cytoplasmic dilution

Previous computational studies have suggested that the two positive feedback loops formed by Cdk1 with its activating phosphatase Cdc25 and inhibitory kinase Wee1 underlie the robustness of mitotic oscillations against molecular noise and parameter perturbations (16,23). To explore whether these Cdk1 positive feedbacks contribute to the cell cycle robustness to cytoplasmic density changes, we performed dilution experiments as before but in the presence of Wee1 inhibitors (0 -5 μM, PD166285) or Cdc25 inhibitors (0 -50 μM, NSC95397). We found that diminishing Wee1 or Cdc25 activity had no noticeable effect on the ability of the system to oscillate (Fig 4A and 4B). As summarized in Fig 4C (Wee1 inhibition) and Fig 4D (Cdc25 inhibition), the system maintained a 100% oscillation percentage (Top panels) and stable period (Bottom panels) for low to moderate dilutions regardless of the concentrations of inhibitors, suggesting the compromised positive feedbacks do not affect much the cell cycle robustness to cytoplasmic density dilution. Nevertheless, Wee1 and Cdc25 inhibitions could modify the period itself independent of dilution, consistent with established knowledge of their role as part of the cell cycle network. Wee1 inhibition causes the threshold for mitotic entry to be lower (less cyclin B production is needed) and thus shortens the period (Fig 4C Bottom Inset, Fig S7); on the contrary, inhibiting Cdc25 increases the amount of cyclin B needed for mitotic entry, which causes the period to elongate (Fig 4D Bottom Inset, Fig S8). These trends agree with our model simulated results (Fig S9, SI text S4). These investigations suggest fine-tuning of the feedback motifs does not appear essential to sustain cell cycle robustness to changes in cytoplasmic density.

**Figure 4.**
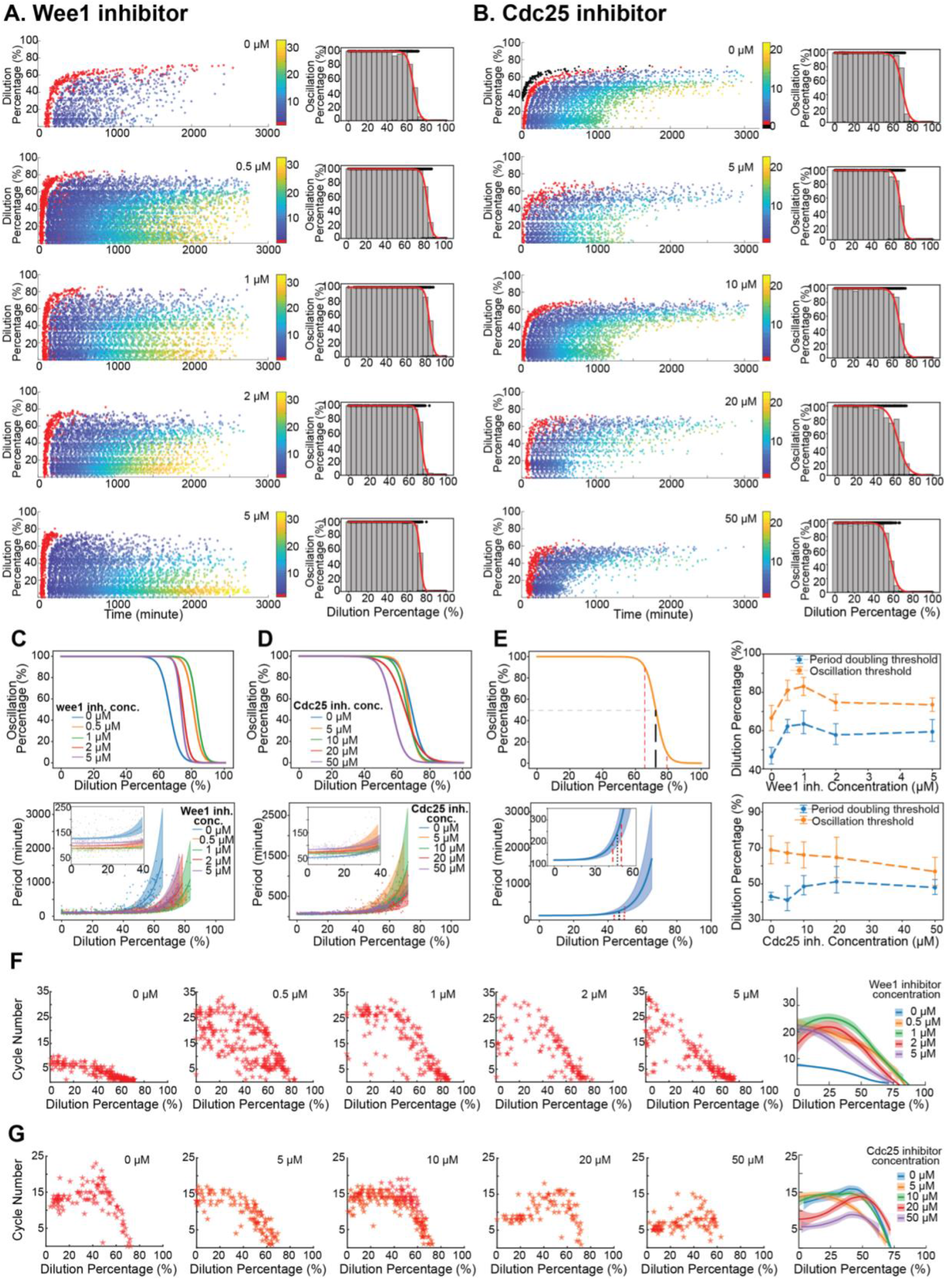
Cdk1 positive feedbacks are not essential for robustness to cytoplasmic dilution. **(A, B)** Extracts were supplemented with 0 μM, 0.5 μM, 1 μM, 2 μM, and 5 μM of Wee1 activity’s inhibitor PD166285 (A), or with 0 μM, 5 μM, 10 μM, 20 μM, and 50 μM NSC95397 for inhibiting Cdc25 phosphatase activity (B), and then tested for their robustness to dilution. For each inhibitor concentration, the raster plot of the experimental result is shown as well as the oscillation percentage obtained. The system retains its function for most of the dilution range (0% to 60%) for all inhibitor concentrations with slight difference in the absolute threshold values and the onset time and end time of oscillations, indicating cell cycle remains robust to dilution despite the disruption of Wee1/Cdc25 positive feedbacks. **(C, D)** Top panels show all oscillation percentage curves together to highlight the differences observed near the thresholds at different inhibitor concentrations of PD166285 (C) and NSC95397 (D). Bottom panels show the obtained period profiles as a function of dilutions. Insets are zoom-in views for the changes in period for low dilutions where the curve is mostly flat. The addition of wee1 inhibitors accelerates cell cycle period as undiluted extracts without wee1 inhibitors show the longest periods compared to other undiluted extracts with inhibitors. The period lengthening effect of Cdc25 inhibitor is significant if we compare undiluted (or low diluted) extracts with different inhibitor levels. Period curves were fitted to exponential functions and bootstrap was performed on the data to obtain confidence intervals for the parameters. Shaded region represents one standard deviation from the average parameters. Quantification of robustness to dilution. Left panels show two different ways of measuring robustness based on oscillation percentage (Top) and period curves (Bottom). For oscillation percentage we extract the dilution required for 90%, 50%, and 10% oscillation percentage, and define the oscillation threshold as the dilution percentage at 50% oscillation percentage. For period curves we calculate the dilution percentage at which period doubles, to be defined as the period-doubling threshold. The estimated error on the parameters is used to infer a confidence interval for the period-doubling dilution. Right panels show how the oscillation and period doubling threshold change as a function of inhibitor concentration. Top panel corresponds to Wee1 inhibitor experiments and the bottom panel to Cdc25. **(F, G)** Total number of cycles as a function of dilution percentage for different concentrations of Wee1 inhibitor (F) and Cdc25 inhibitor (G). Rightmost plots show all the curves together smoothed by locally weighted scatterplot smoothing (lowess) for ease of comparison. It can be appreciated that the inhibition of Wee1 increases the cycle number for low dilutions. The addition of the Cdc25 inhibitor tends to decrease the number of cycles in the system for most dilutions. Notably, in (A, C, E, F), cell cycle robustness and cycle numbers in response to dilution show a non-monotonic behavior with respect to Wee1 inhibitor concentrations, suggesting a peak performance (with highest robustness and largest number of cycles) at an intermediate 1 μM Wee1 inhibitor. In contrast, the system’s response to Cdc25 modulation is monotonic

On the other hand, the system’s responses at high dilution (>60%) showed a dependency on positive feedback inhibitions. The half-maximal threshold for the oscillation percentage increased and then decreased as the Wee1 inhibitor concentration increased, yielding a non-monotonic function of Wee1 inhibition (Fig 4C, Top). On the contrary, adding Cdc25 inhibitor seemed to monotonously decrease the oscillation threshold (Fig 4D, Top). To quantify the qualitative changes observed in both oscillation percentage and period of the system, we fit the data curves with logistic regression and an exponential function, respectively. These fits allowed us to extract the half-maximal threshold (Fig 4E Top Left) as well as the dilution percentage value at which the period doubles from its value at 0% dilution (defined as the period-doubling threshold, Fig 4E Bottom Left). We observed that both half-maximal and period-doubling thresholds followed a non-monotonic trend when the Wee1 inhibitor was added (Fig 4E Top Right). On the other hand, adding Cdc25 inhibitor decreased the half-maximal threshold monotonously while the period-doubling value increased slightly. For both cases, we observed similar trends in simulations (Fig S9, SI text S4).

Interestingly, the total number of cycles showed notable changes with the Cdk1 positive feedback inhibitions. Wee1 inhibition increased the total number of cycles from a maximum of 10 (Wee1 inhibitor of 0 μM) up to 33 cycles in maximum (Wee1 inhibitor of 5 μM) (Fig 4F). The system with Wee1 inhibition also presented a longer overall oscillation time than the system without (Fig 4A). In contrast, Cdc25 inhibition reduced the system’s ability to sustain oscillations for many cycles and a long time. As the concentration of Cdc25 inhibitor rose, the total number of cycles decreased (Fig 4G), and the overall oscillation time was shortened (Fig 4B). In summary, we find that both dilution and feedback strength modify the total number of cycles in the system, suggesting a previously unexplored role of Wee1 and Cdc25 in modulating the cycle number in oscillating extracts.

## DISCUSSION

We have investigated how the variations in cytoplasmic density affect cell cycle functioning. Our experiments *in vitro* imply that the cell cycle network can carry out oscillations invariant to a large span of changes above or below the density of an endogenous cytoplasm of *Xenopus* eggs. On the other hand, near the boundaries of the physiological range (0.2-1.22x RCD), the system becomes sensitive to cytoplasmic density variation and switches from oscillation to an arrested state in a sharp, sigmoidal manner, revealing a role of the cytoplasm in controlling the states of a cell when too diluted or concentrated.

Studies have also suggested density of cytoplasm is involved in cell homeostasis. In fission yeast, cells with higher cytoplasmic density (higher total protein concentration) tend to undergo supergrowth at a higher rate to achieve proteome homeostasis (24). In human cells, cytoplasmic dilution of the cell cycle inhibitor Rb through cell growth in the G1 phase has been shown to trigger cell division, providing a mechanism to promote cell size homeostasis (25). Interestingly, while the cell cycle period is highly tunable to the cyclin B mRNA concentration (varied by adding recombinant mRNAs or morpholinos of cyclin B) (26), we found their period is very robust to the density changes of the whole cytoplasm even though this also causes variations in the cytoplasmic cyclin B mRNA concentration. We hypothesized that instead of the absolute concentrations of each component, ratios between components are what influence the robustness of the mitotic oscillator. This divergent response of cell cycles to changes in global concentration versus individual molecular species concentration may help decouple possible cell homeostasis control mechanisms from cell cycle progression.

We also demonstrate that the concentration-induced cell cycle arrest can be reversed by dilution. This reversible phenomenon rules out the possibility of apoptosis, which could also be associated with cell shrinkage and condensed cytoplasm (27), and indicates the existence of other mechanisms to stop the cell cycle. Furthermore, we find that bringing an arrested extract to oscillate via dilution requires a significantly lower density (0.87x RCD) than that for arresting the extract through concentration (between 1.22x and 1.46x RCD). This threshold difference may come from the dynamical properties of the cell cycle interaction network, as suggested by mathematical modeling. A hysteresis produced by a subcritical Hopf bifurcation causes different thresholds to switch between the oscillation/non-oscillation states depending on which state the system originates. The existence of this type of bifurcation has also been reported theoretically in the cell cycle (28) and other biological systems such as *B. Subtilis* biofilms (29). To our knowledge, the present work provides the first experimental observation of a dynamical hysteresis in mitotic oscillations. Moreover, multiple studies have previously analyzed the hysteresis in switches between two stable steady-states of CDK1 in response to the total concentration of its regulatory protein cyclin B. Here, in contrast, we show that oscillatory behavior presents hysteresis as a function of cytoplasmic density. This opens the possibility to the presence of hysteresis with respect to other parameters of the system, for example, cyclin B synthesis rate, rather than the total protein of cyclin B in case of bistable switch of CDK1. Future experiments could test whether oscillations appear and disappear for the same threshold value when increasing and decreasing the synthesis rate of cyclin B.

Hysteresis is widely present in physics, chemistry, and biology. Our discovery of the hysteretic response of the cell cycle to cytoplasmic density adds to a new example of this important universal phenomenon. It is not yet clear the advantage for the cell cycle to possess this property, but hysteresis could function as a memory switch for cells that have undergone extreme cytoplasmic concentrations and entered an altered state, so they do not easily return to normal cell cycle progression by small fluctuations. Curiously, the model does not predict the same level of hysteresis to happen for the other end of extremes, that is, to arrest the system via dilution and recover its oscillations by concentration. Further experimentation could investigate this effect by concentrating highly diluted extracts and comparing the observed thresholds.

Our model could sufficiently reproduce most experimental observations by assuming that cyclin B synthesis rate, degradation rate, and protein’s total concentration depend on cytoplasmic density. Interestingly, Neurohr et al. find that in oversized cells (cell density reduction from ∼1.10 to ∼1.07) (9), transcription and translation machinery become limiting and do not scale with cell size, suggesting that our assumption of decreased synthesis rate with cytoplasmic density might be appropriate. However, more detailed modeling of these fundamental processes is needed to understand the mechanisms by which synthesis and degradation are affected. Quantitative cell-free experimentation with minimal systems and bottom-up approaches are well-suited to examine the role of cytoplasmic density on protein synthesis and degradation in future studies. Meanwhile, our coarse-grained model does not capture every dynamical observation of the system. For example, when concentrated, extracts settle into a high CDK1 activity level while our model settles to a relatively low activity level. The high CDK1 activity could be explained by accumulated cyclin B for highly concentrated extracts that increase protein synthesis rate. However, our model does not explicitly include the dynamics of cyclin B binding and unbinding with CDK1. Therefore, it prevents such accumulation and eventual saturation of CDK1 molecules. An expansion of our current model could help in underpinning the exact mechanisms by which this network achieves robustness when cytoplasmic density is modified.

Robustness to changes in environmental variables has been observed in other biological oscillators. Studies have shown neuronal oscillations are robust to changes in temperature and pH while at the same time displaying a rich dynamical structure (30). Similarly, the KaiABC circadian oscillator remains unchanged when the concentration of all components is scaled at constant ATP initial input (from 5.0x to 0.1x) but settles into a steady-state when further diluted (31).

We demonstrate that modifying the strength of Cdk1’s positive feedback does not compromise the robustness to changes in cytoplasmic density. This implies that fine-tuning of these interactions are not essential for the robustness to dilution. What interactions of the network are responsible for robustness to changes in cytoplasmic density are still unknown. A sensitivity analysis of our model suggests various candidates for future experimentation (Fig S10, SI text S5). In particular, both PP2A and Greatwall associated reactions appear as promising candidates. On the flip side, Wee1 and Cdc25 inhibition modified the total amount of cycles observed in extracts. Wee1 inhibition increased the total number of cycles while Cdc25 inhibition decreased them. This novel observation suggests that the termination of the cell cycle in cell-free cytoplasm is tightly regulated, and might not be entirely due to the total consumption of resources such as energy as previously speculated (17). This opens the possibility to consider several hypotheses for the stopping of oscillations. First, extracts could recapitulate certain events associated with midblastula transition (MBT) in normal development. Namely, maternal mRNA is cleared and checkpoints arise in the cell cycle (32). Since no DNA is present in our extract preparations, mRNA clearance could mean the depletion of cyclin B mRNA from the system, and thus elongation of the cell cycle and eventually termination. In fact, RNA-seq studies show that cyclin B mRNA is depleted after stage 12 of *Xenopus laevis* development (33). Real-time PCR could aid in the investigation of this hypothesis. Also, it could be possible that checkpoints arise within the extract in a timed fashion. The main checkpoint arising after MBT is mediated by the protein Chk1, which shares a signaling pathway with Wee1 and Cdc25 via the protein 14-3-3 (34,35). Tracking Chk1 concentration and activity throughout the lifespan of an extract could shed light on this hypothesis. In either case, our results present a new avenue of research for understanding what regulates the total number of cycles in *Xenopus* egg extracts.

Finally, our results could be expanded by including nuclei in our extract preparations, which could be achieved by supplying demembranated sperm chromatin into *Xenopus* extracts. Theoretical work suggested that the extra dynamic bistable switch induced by nuclear translocation promotes robustness of cell cycle oscillations to noise (36). How changes in cytoplasmic density affect a system where nuclei are present is currently unknown and can be a promising avenue for research. It has been reported that the relative density of the nucleus and cytoplasm is robust during cell growth in human cells (37), which opens the question of whether a balance between the density of the nucleus and the cytoplasm is necessary for the correct progression of the cell cycle. Importantly, the spatial self-organization in *Xenopus* egg extracts has also been shown to be robust to dilution (38), expanding our results from the time domain to the spatial domain. How both spatial and temporal dynamics together are affected by dilution appears as an exciting new direction of research. The experimental platform developed in the current study can be easily generalizable to quantitatively studying various other cellular processes in density-varied cytoplasm.

## MATERIALS AND METHODS

### Cycling Xenopus laevis extract preparation

Cycling Xenopus extracts were prepared following previous protocol with a few modifications (17). Sexually mature female frogs were purchased from Xenopus1 and Nasco Education. We primed frogs for ovulation with an injection of 100 IU of human chorionic gonadotropin (HCG) (02198591, MP Biomedicals) one week before and induced them with 600 IU of HCG 12-16 hours before acquiring the eggs. The eggs were activated by 0.5 μg/ml calcium ionophore A23187 (C7522, MilliporeSigma). A 10-minute followed by a 5-minute centrifugation at 20,000g was conducted in a Beckman Avanti J-E Centrifuge with a JS-13.1 swinging bucket rotor to crush eggs and extract cytosolic materials. ATP-mix was not added. In dilution experiments, extracts were diluted with freshly prepared Extract Buffer (100 mM KCl, 1 mM MgCl2, 0.1 mM CaCl2). In concentrating experiments, bulk extracts were loaded into a 96-well microplate with 40 μl per well, and the whole plate was placed in a vacuum for 10-minute intervals up to 40 minutes. After each time interval, concentrated extracts in two or three wells were taken out and combined to ensure a minimum sample volume of 50 μl for each condition. The concentrating efficiency was estimated by the changes in Nanodrop measurement, volume loss, and Alexa Fluor 594 dextran intensity before and after evaporation (details shown in Table S1). Notably, the average intensity of droplets after evaporation was calculated by fitting a normal distribution to the histogram of droplet intensity.

### Fluorescence-labeled reporters and inhibitors

CDK1-FRET sensors were prepared following a method described earlier (G.M. and Q.Y., unpublished work). A final concentration of 1μM CDK1-FRET sensors was added to monitor the dynamics of CDK1/PP2A activities. Dextran-Alexa Fluor 594 (D22913, Invitrogen) at a final concentration of 200 nM was added to extracts for measuring the dilution percentage of extracts encapsulated within each droplet. For experiments shown in Fig 4, 0-5 μM of PD166285 (PZ0116, MilliporeSigma), or 0-50 μM of NSC95397 (N1786, MilliporeSigma) was added into bulk extracts to manipulate positive feedback strengths.

### Device fabrication, tuning, and droplet generation

Two-channel microfluidic device fabrication, droplet generation, and droplet loading were performed as previously reported (20). Dow Sylgard 184 silicone encapsulant (4019862, Ellsworth) was poured on SU-8 modes on silica wafers to produce PDMS slabs with microfluidic channels. In a 2-channel microfluidic device, extracts supplied with dextran-Alexa Fluor 594 and CDK1-FRET sensor were flown in from one inlet via a transfer tubing (EW-0641711, Cole-Parmer), while Extract Buffer for dilution with the same concentration of CDK1-FRET sensor was flown in from another inlet. A multichannel pressure controller (OB1 MK3+, Elveflow) controlled the flow rate in each inlet precisely. By changing the temporal pressure profiles of individual inlets periodically while keeping the total aqueous pressure constant, we mixed extracts with the extract buffer at different ratios, covering the whole 0-100% dilution spectrum continuously. The boundaries of 0% and 100% dilution were confirmed by the observation of one aqueous flow pushing the other back to its inlet completely. These diluted extracts were then encapsulated by 2 weight % 008-FluoroSurfactant oil (008-FluoroSurfactant-2wtH-50g, Ran Biotechnologies) into uniform-sized droplets. The size of droplets examined in most experiments has a diameter between 70 μm to 90 μm, except the smaller droplets (55.29±1.83 μm in diameter) investigated in Fig S2. The generated droplets were collected in the reservoir of the microfluidic device, and via a simple dipping step, these droplets were loaded into hollow glass tubes (inner height: 100 μm; width: 2 mm, 5012, VitroCom) to form a single layer of droplets. These glass tubes were pre-cut into 3-5 mm pieces, and precoated with Trichloro (1H,1H,2H,2H-perfluorooctyl)silane (448931, MilliporeSigma). Loaded glass tubes were then placed in a glass-bottom sterile Petri dish (70674, Electron Microscopy Sciences) and immersed with mineral oil (8012951, Macron Fine Chemicals) to prevent surfactant oil and sample evaporation.

### Time-lapse fluorescence microscopy and image processing

All imaging was carried out at room temperature on an inverted Olympus IX83 fluorescence microscope with a 4x air objective, a LED fluorescence light source, a motorized x-y stage, and a digital CMOS camera (C13440-20CU, Hamamatsu). The open source software μManager v1.4.23 was used to control the automated imaging acquisition. Bright-field and multiple fluorescence images (CFP/FRET/RFP) of samples were recorded at a frequency of 1 cycle/4-8 minutes for 1-3 days. Custom scripts in MATLAB 2019a were written to perform image processing. Bright-field images were used for individual droplet segmentation and tracking. Only individual droplets tracked from the beginning (first 50 frames) were selected for analysis. Intensity peaks and troughs were first auto-selected by MATLAB and then manually corrected for reliability. Other oscillation characteristics, like rising periods and falling periods, were also calculated for further analysis.

### Statistical analysis

In dilution experiments, Alexa Fluor 594 intensities were used as references to quantify dilution percentage. The RFP intensity of individual droplets were first sorted and normalized, with the 99th percentile defined as 0% dilution and the 1st percentile defined as 100% dilution. For Fig 1E, 1F, 4A, 4B, 4C top, and 4D top, logistic regression was employed to infer the percentage of oscillation as a function of dilution. For each experiment, individual droplets were recorded as either oscillatory or non-oscillatory, and this discrete data was used for the regression. Fitting was performed in Python 3.7.10 using the logistic regression function from the package scikit-learn 0.22.2. Each bar in Fig 1E, 2E,4A, and 4B represents a percentage of oscillatory droplets over total droplets detected within the bin range. Fig 2E applied nonlinear least-square fitting from the function curve_fit to the data of concentrating extracts and the dilution of concentrated extracts. The fitting results were shown in orange and blue, respectively. In Fig 3C, 3G, and 3J, simulation results were directly transformed to oscillation percentage by taking the ratio of oscillatory droplets and total simulated systems. For comparison with experimental curves, a logistic function was obtained from the simulation results via fitting using the function curve_fit from the package scipy 1.4.1. The same functionality was used to fit an exponential function to the period vs. dilution curves presented in 4C bottom and 4D bottom. Confidence intervals are obtained by bootstrapping the data. Specifically, data was resampled (the new sample size was the same as the original sample size) with replacement 104 times. For each resampling, the new set of points was fitted and the coefficients recorded. 25th and 75th percentiles of the parameter distributions were taken as estimates on the uncertainty of the parameters. Shaded areas in 4C bottom and 4D bottom represent the obtained curves with the limit uncertainty values of each parameter. In Fig 4E and 4F, locally weighted scatterplot smoothing (LOWESS) with a span of 0.75 was applied to smooth the total cycle number curves for different inhibitor concentrations. Package ggplot2 in Rstudio 1.2.5019 was installed and utilized. Shaded areas represent the moving first and third quantiles.

### Mathematical modeling

All time series were obtained by numerical integration of the ODE equations. Simulations were performed using custom scripts in Python 3.7.10. Oscillatory or steady-state behavior was differentiated by the norm of the system’s velocity-vector in conjunction with an analysis of the signal’s peak detection features. Droplet simulations are ODE simulations where the corresponding total concentrations, synthesis rate, and degradation rate were sampled from a Gamma distribution representing the encapsulation error. In all cases, 500 droplets were simulated per RCD value. The bifurcation diagram in 3I was performed in XPPAUT 8.0. The diagram was started from a high dilution converged steady state. Stable and unstable steady states were first tracked while the Hopf bifurcations were continued in a second iteration of the algorithm. Model equations and setup details were provided in the accompanying SI texts.

## ACKNOWLEDGEMENTS

We thank Gembu Maryu for CDK1-FRET sensor construction and preparation, Meng Sun for microfluidics training, Zhengda Li for help with image processing code troubleshooting, and all members in Yang Lab who contributed to the discussion of this project. This work was supported by NSF (MCB #1817909; Early Career #1553031), NIH (NIGMS #R35GM119688), and Alfred P. Sloan Foundation.

## Author Contributions

Designed research (MJ, FT, SW, QY), performed experiments (MJ), developed model (FT), analyzed results (MJ, FT, QY), wrote manuscript (FT, MJ, QY).

## SUPPLEMENTARY TABLES, FIGURES, AND VIDEOS

**Table S1:**
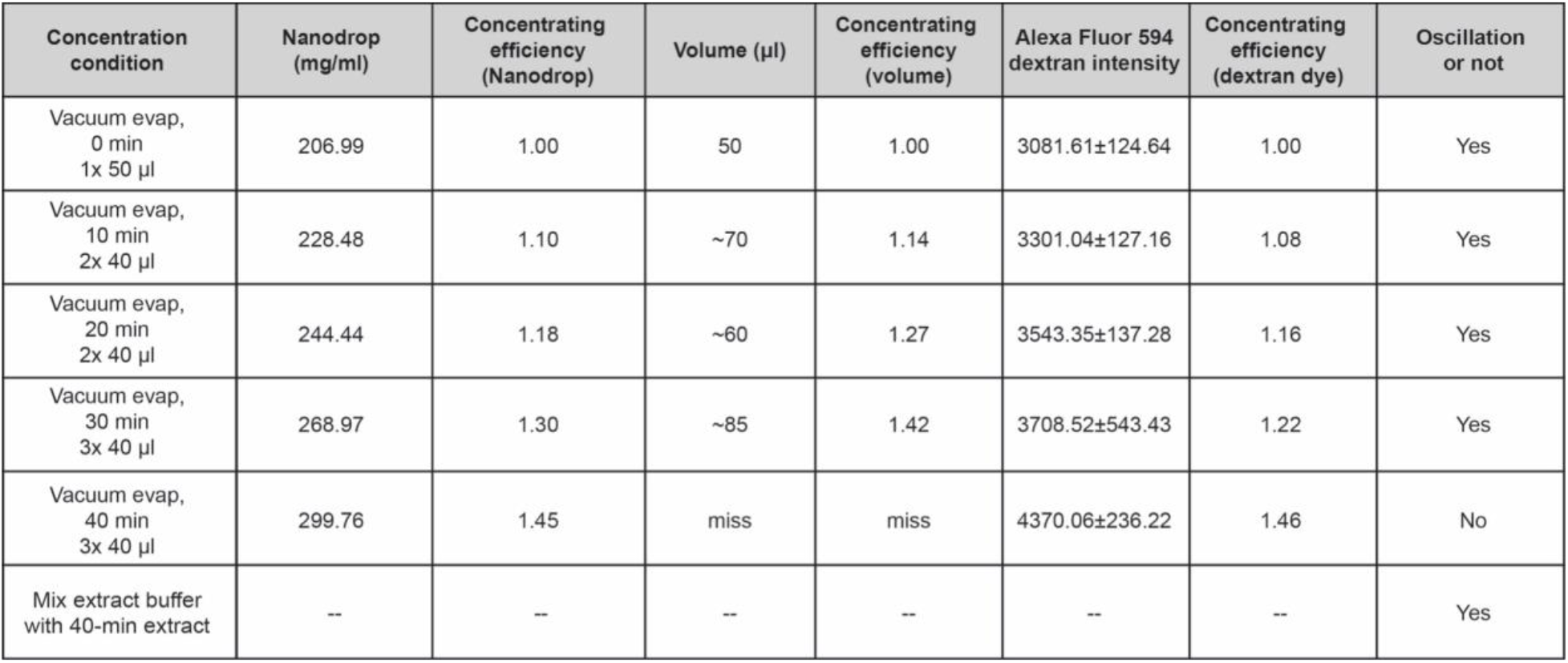
Three methods are applied to calculate the concentrating efficiency of vacuum evaporation. Dextran intensity of each condition is quantified from fitting the histogram of droplet intensity to a normal distribution. By comparing these three methods, we choose fluorescence intensity changes before and after evaporation as the primary method for quantification.

**Figure S1:**
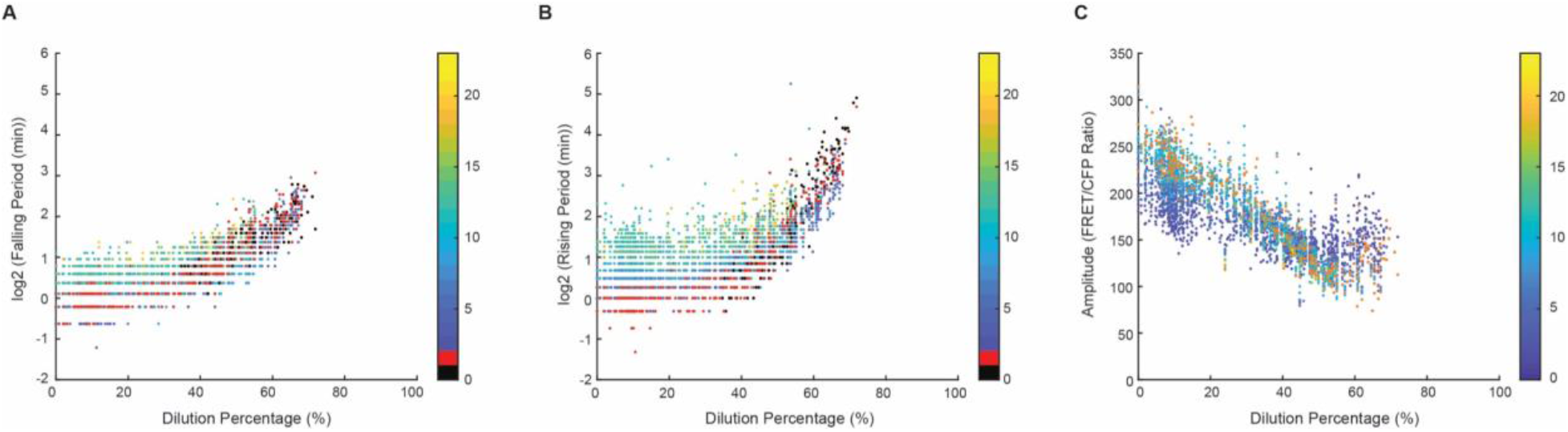
Dilution influences rising periods more pronouncedly than falling periods. (A) Fold changes of rising period (interphase) and (B) falling period (mitosis) for the extract in Fig 1. Dots in black are cycle 0 and dots in red are associated with cycle 1. Both interphase and mitosis are robust up to 40% dilution percentage (0.6x RCD), whereas rising periods have more significant fold changes (up to 4.8x) compared to falling periods (up to 2.9x) at high dilutions. Thus, interphase is more impacted than mitosis by dilution, manifesting the tighter regulations during mitosis. (C) the dynamic of amplitude as a function of dilution for the extract in Fig 1. Each dot represents one cycle. Amplitudes of the last cycle are usually disrupted and do not show an accurate measurement, so we label them in orange in the plot. Amplitudes first decrease as the system is being diluted, while near the oscillation threshold, amplitudes increase. This increase may be correlated to a significant period elongation in these droplets and the stoichiometry changes between CDK1 and CDK1-FRET sensor in highly diluted extracts.

**Figure S2:**
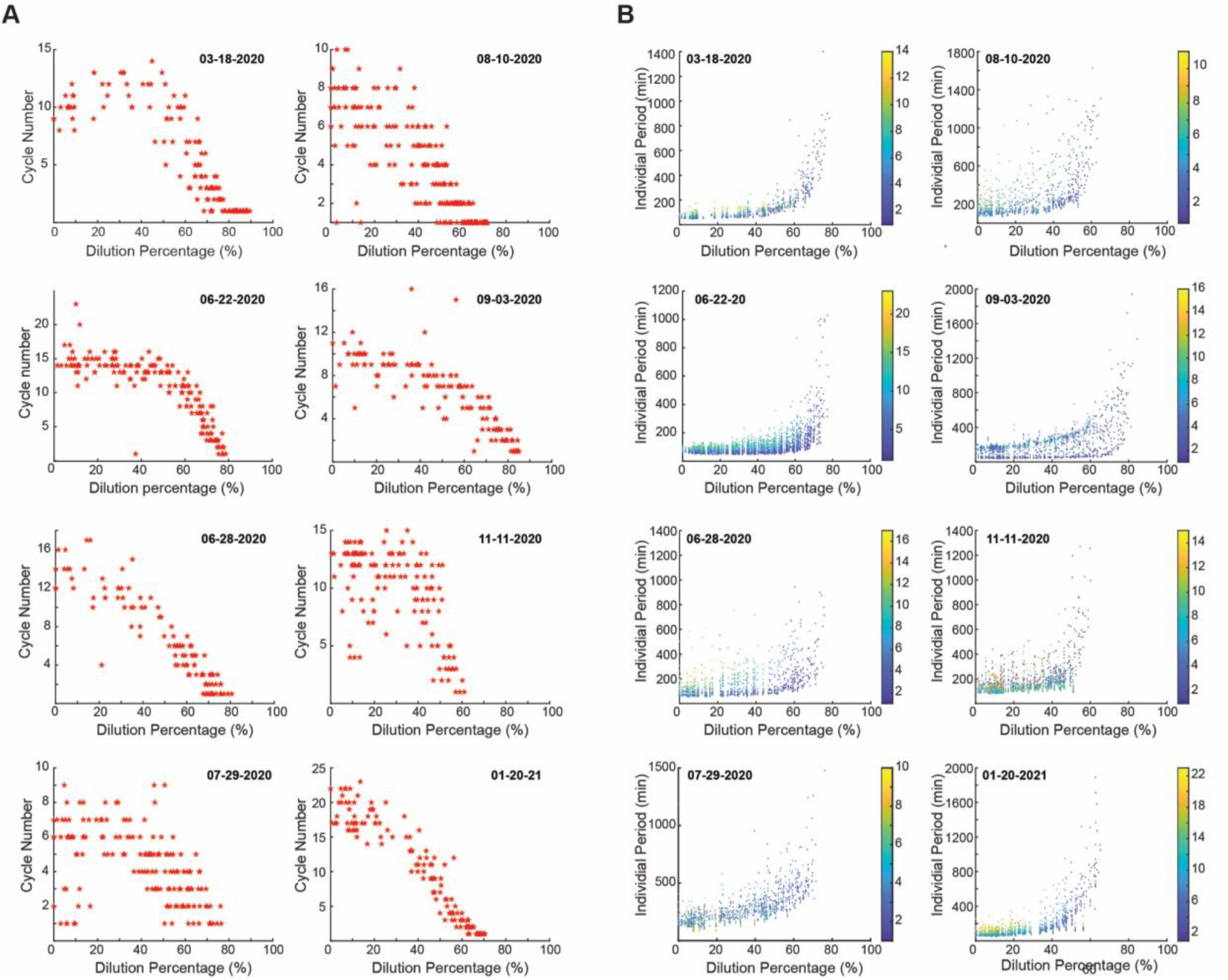
Even though there are some batch-to-batch variations, the overall trends of (A) cycle number and (B) period as a function of dilution are consistent across nine different day preparations (one is shown in Fig 1). Cycle numbers present a slight spike at moderate dilutions in two out of nine experiments; and in other three out of night experiments, cycle numbers decrease relatively linearly with dilutions. In the rest four experiments, cycle numbers show a plateau with low range of dilutions and then decrease abruptly at high dilution percentages. In most experiments, the period curve is relatively flat at the low dilution region and then has a drastic increase as dilution percentage becomes high. Even though the absolute value of where periods start to significantly extend varies across these nine experiments, the overall trend is maintained.

**Figure S3:**
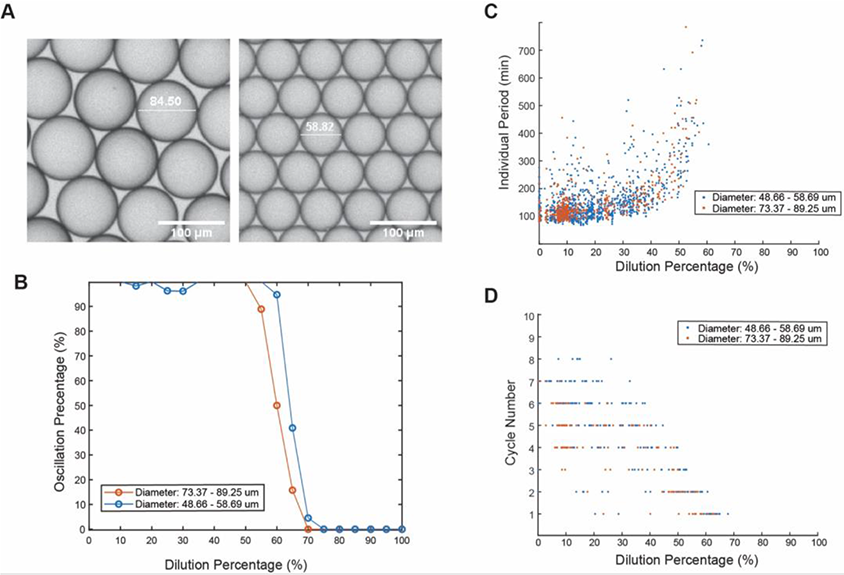
Partition errors are not associated with the halt of oscillation in highly diluted droplets. Droplets with (A) two different diameters: 83.18±2.21 μm and 55.29±1.83 μm show no significant difference in (B) oscillation percentage, (C) the total period, and (D) cycle number as a function of dilution percentage.

**Figure S4:**
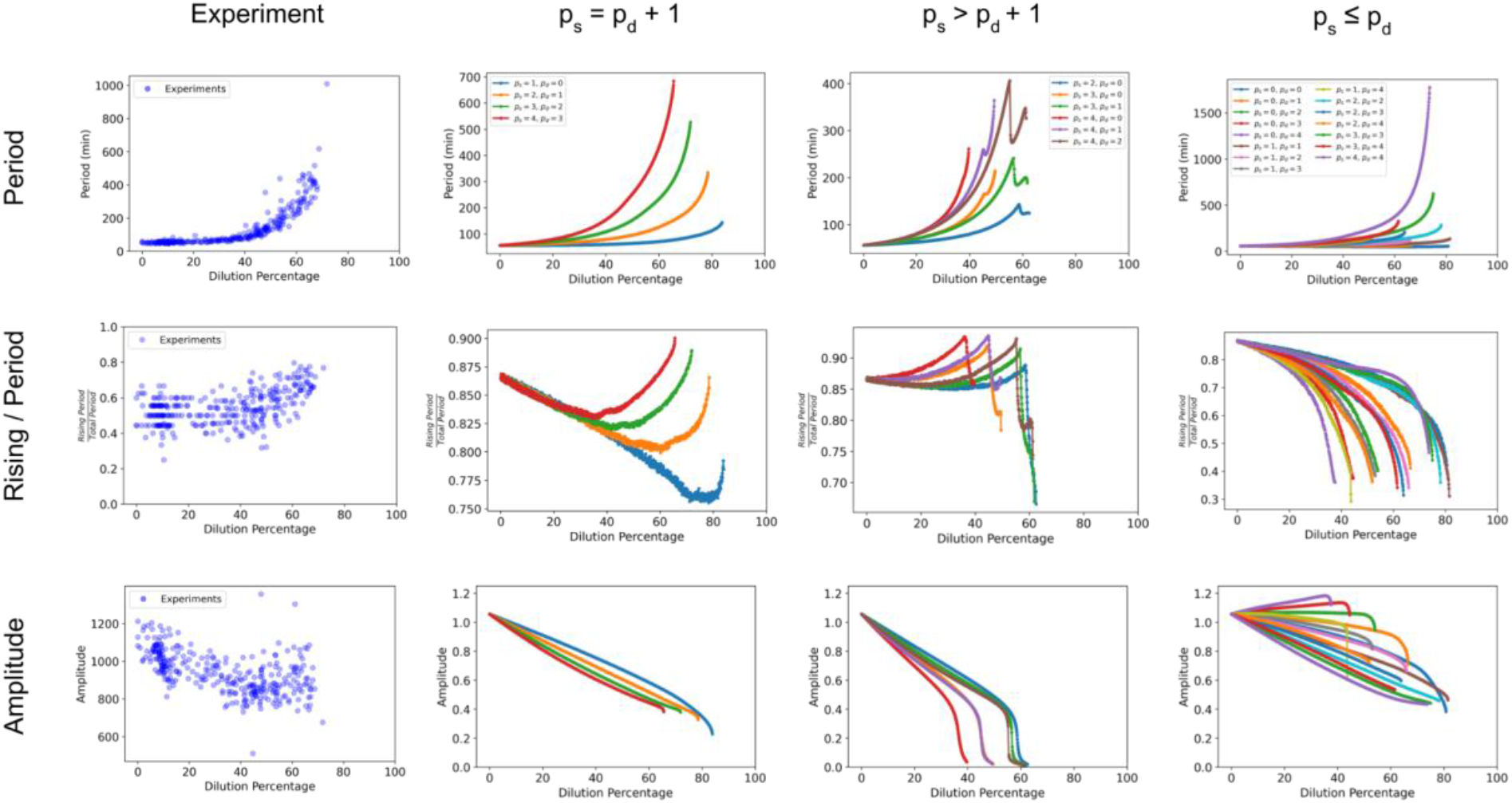
Selection of appropriate dependency of synthesis and degradation rate with cytoplasmic density. **Rows:** Three different quantities are analyzed from experiments and simulations as a function of dilution percentage. The total period, the fraction of the time the system spends in the rising phase, and the amplitude are used to compare experiments with simulations and choose the appropriate scaling for synthesis and degradation rate. **Columns:** Experiments and simulations. Simulations are separated into three groups according to the relationship between ps and pd and the behavior of the three quantities analyzed. All the possible integer combinations of ps and pd between 0 and 4 are studied.

**Figure S5:**
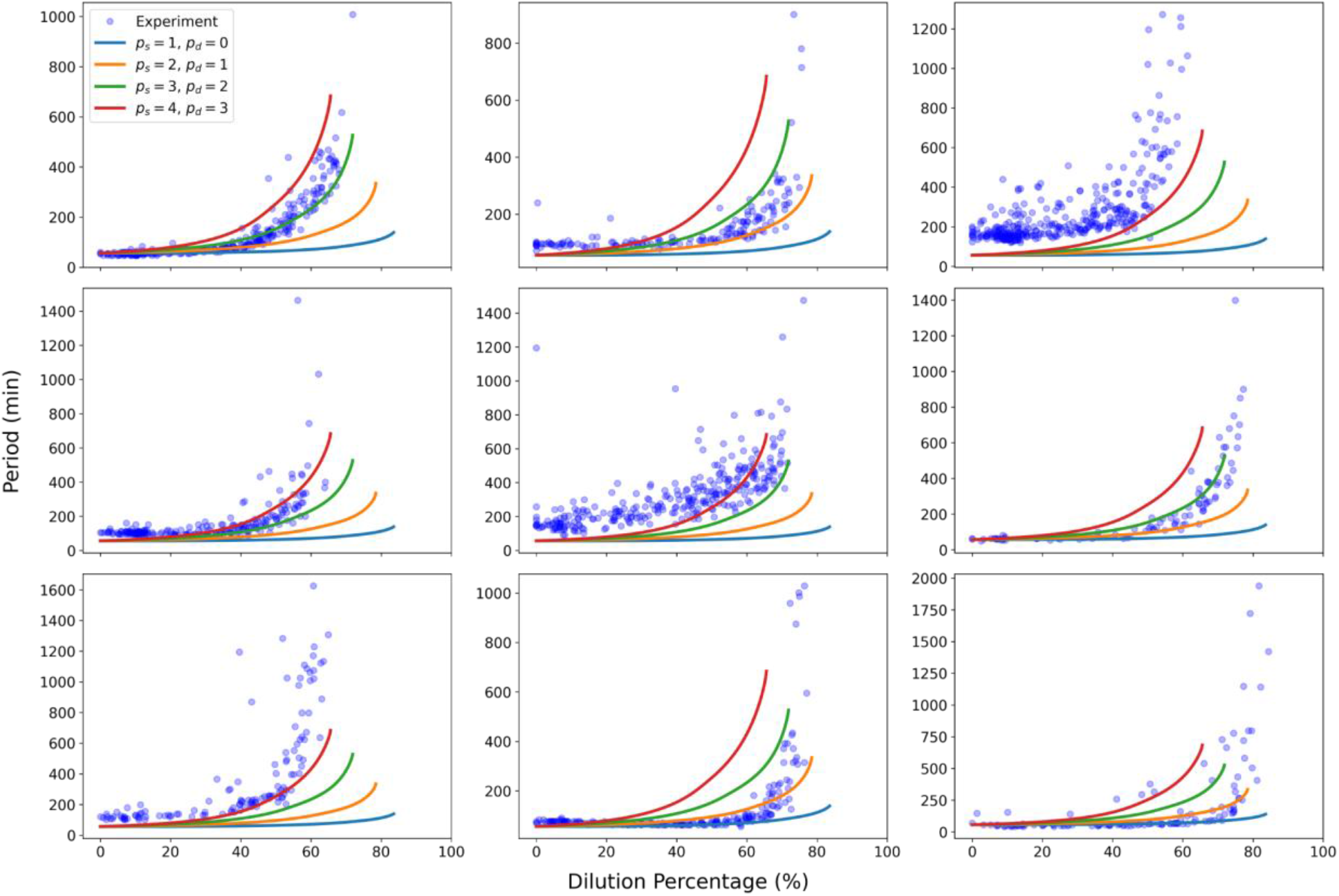
Comparison of selected models with experimental data. Each panel represents a different replicate for the experimental data. In all plots the four selected curves are presented. As a reference the leftmost top plot is the same dataset presented in Fig S4. The case ps = 3, pd = 2 (green curve) was used as a reference for further studies of the model.

**Figure S6:**
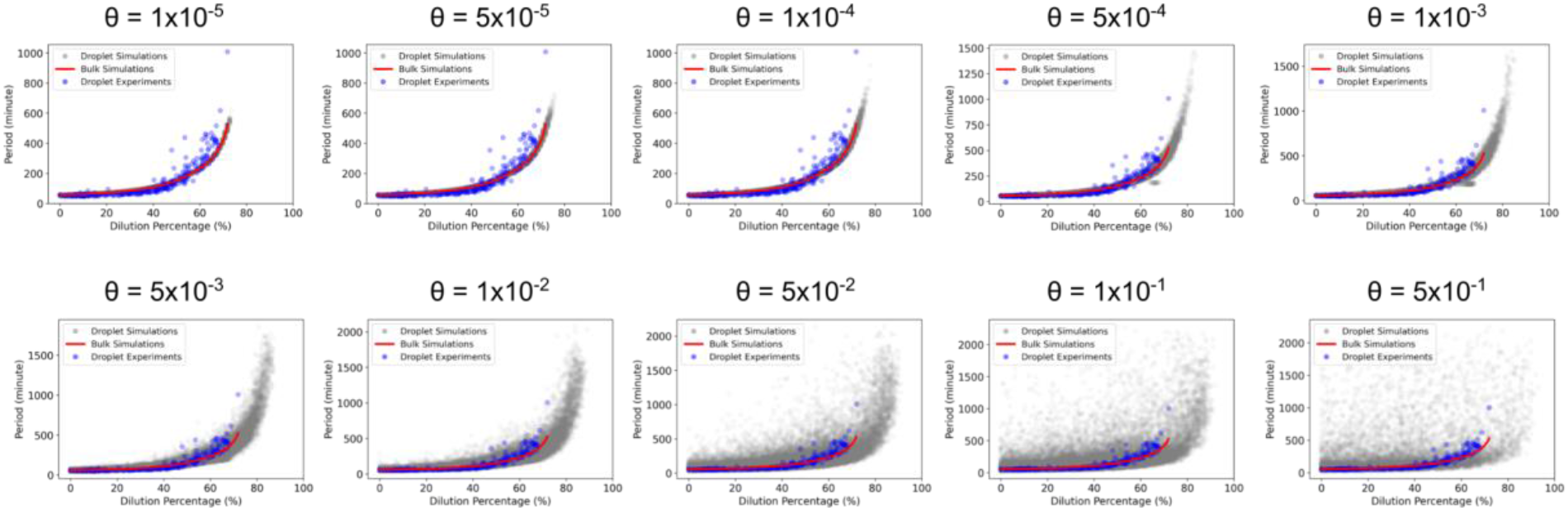
Droplet simulations with different variability in the parameter selection. Theta (θ) values were selected between 10^−5^ (droplets follow closely the bulk solution)and 10^−1^ (droplets deviate significantly from bulk solution). For the main text we selected a value of θ = 10^−4^ as it represented a level of variability similar to the experimental data as well as matched the observed coefficient of variation (CV) of fluorescent probe encapsulation.

**Figure S7:**
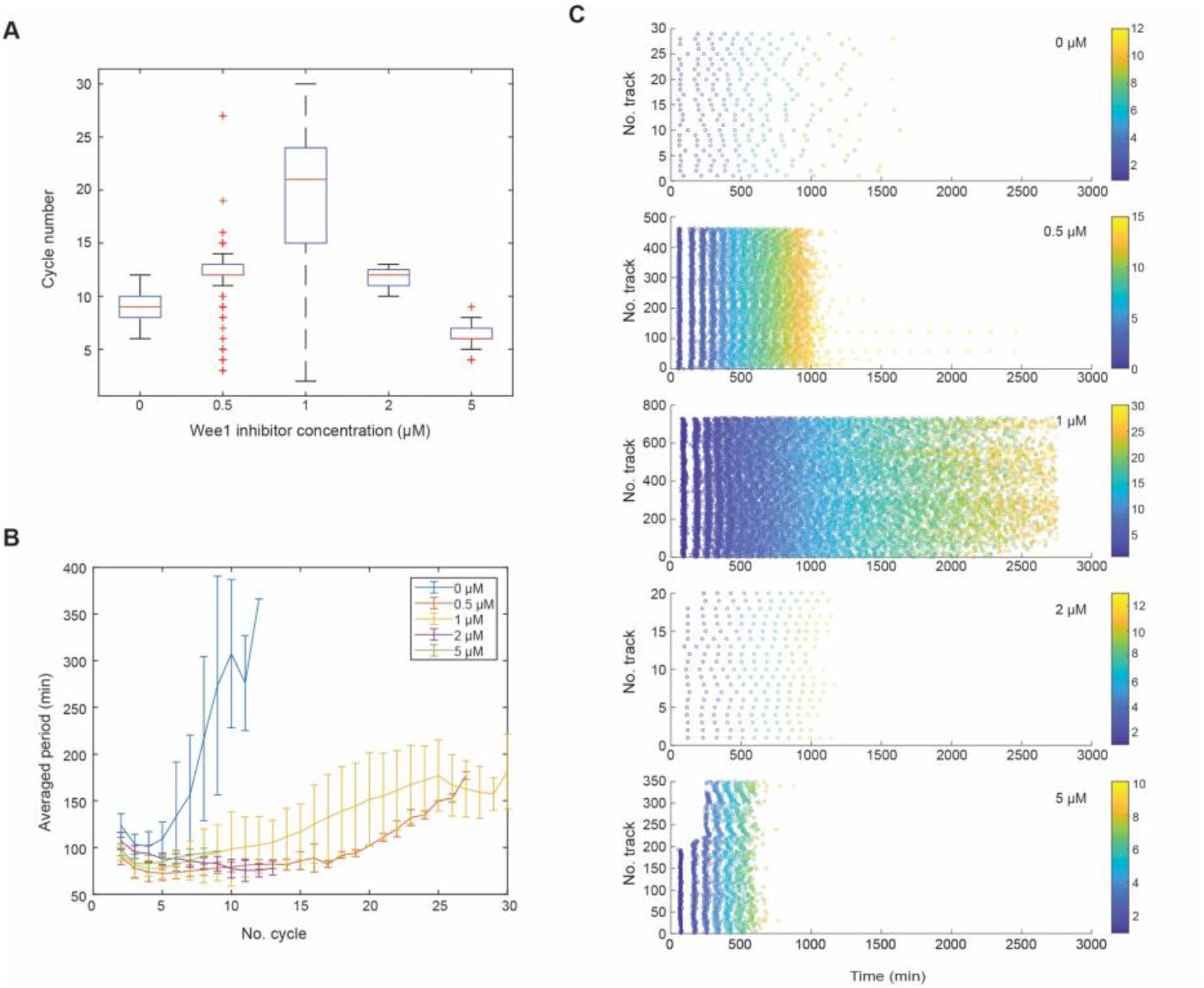
Wee1 inhibitor PD166285 modulates (A) cycle number, (B) the duration of periods, (C) the total oscillation time in extracts without dilutions. The duration of periods is shortened with wee1 inhibitor addition. The cycle number and total oscillation times of the extract respond to the inhibition of wee1 non-monotonically: the system with 1uM wee1 inhibitors shows the maximum cycle numbers and most prolonged total oscillation time.

**Figure S8:**
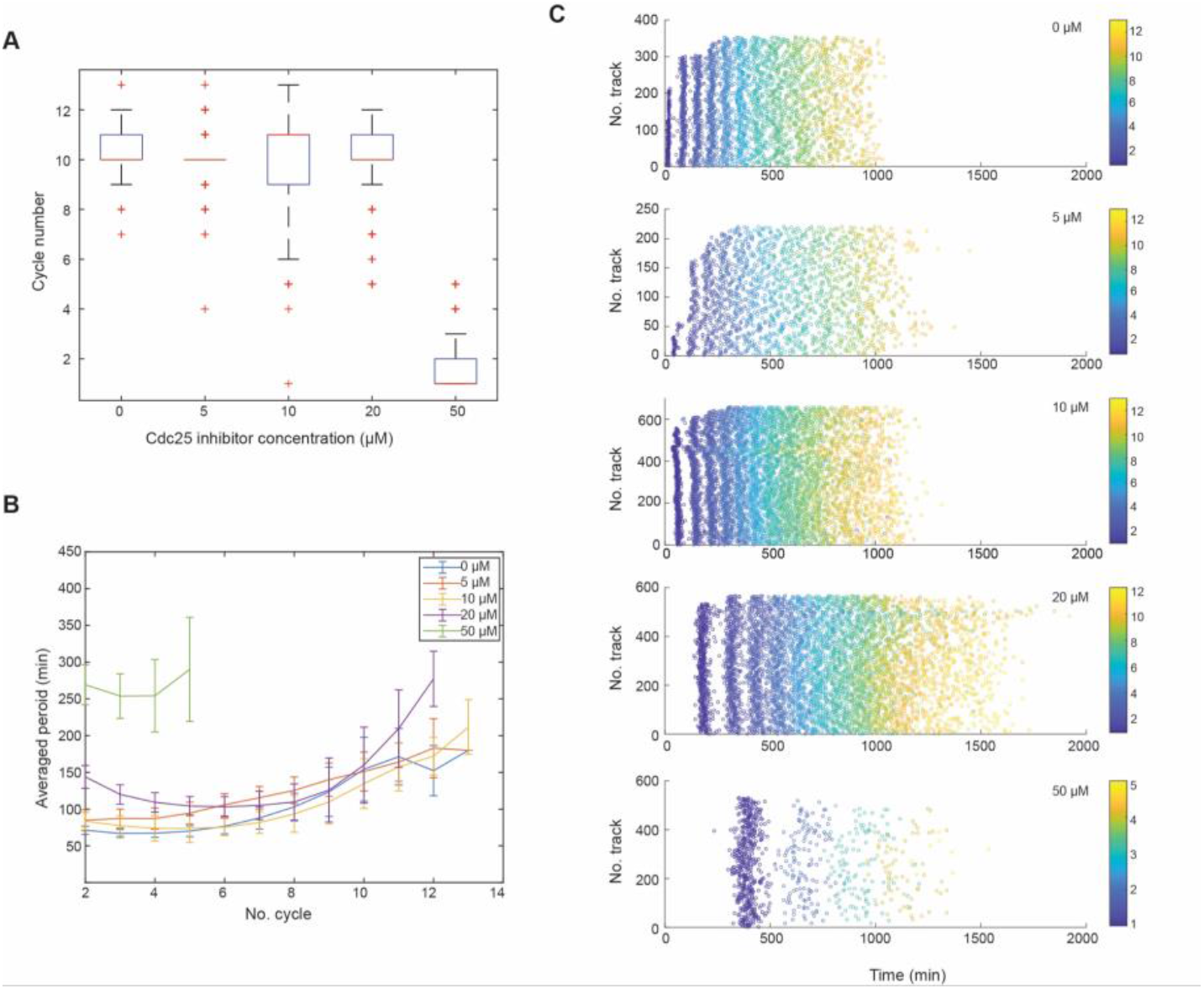
the influence of Cdc25 inhibitor NSC95397 on (A) cycle number, (B) the duration of periods, and (C) the total oscillation time in extracts without dilutions. System with Cdc25 inhibitor addition shows extended period length for early cycles, which is consistent with previous results reported in literature. Extracts with 50 μM inhibitors show less tendency to oscillate, so we assume this inhibition level is too high for maintaining normal oscillation behaviors. Besides the high 50 μM condition, the cycle number does not show a significant correlation with cdc25 inhibition levels, but due to the prolonged period length, the total oscillation time is still extended as cdc25 is being inhibited.

**Figure S9:**
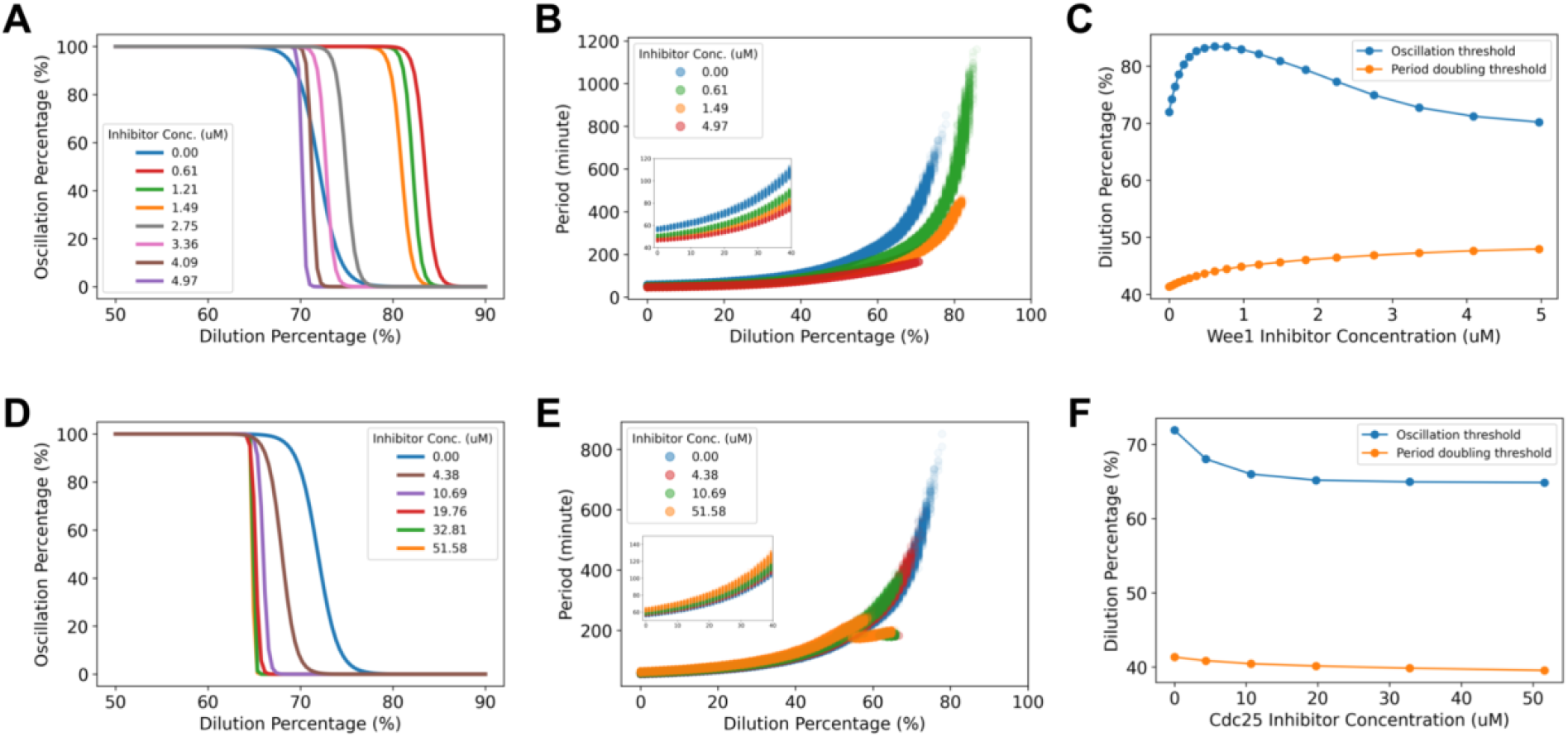
Simulation of the effect of Wee1 and Cdc25 inhibitors on robustness to dilution. For Wee1 inhibition different oscillation percentage curves are shown (A) as well as period (B) and thresholds (C). The same information is shown for Cdc25 inhibition in (D), (E), and (F). Oscillation thresholds are obtained from droplet simulations using the setup described in SI text S6. Period doubling thresholds are obtained from bulk simulations at each corresponding set of parameters. Insets in (B) and (E) show a zoomed version between dilutions 0% and 40%. In (A), (B), (D), and (E) only certain curves are shown for visual clarity.

**Figure S10:**
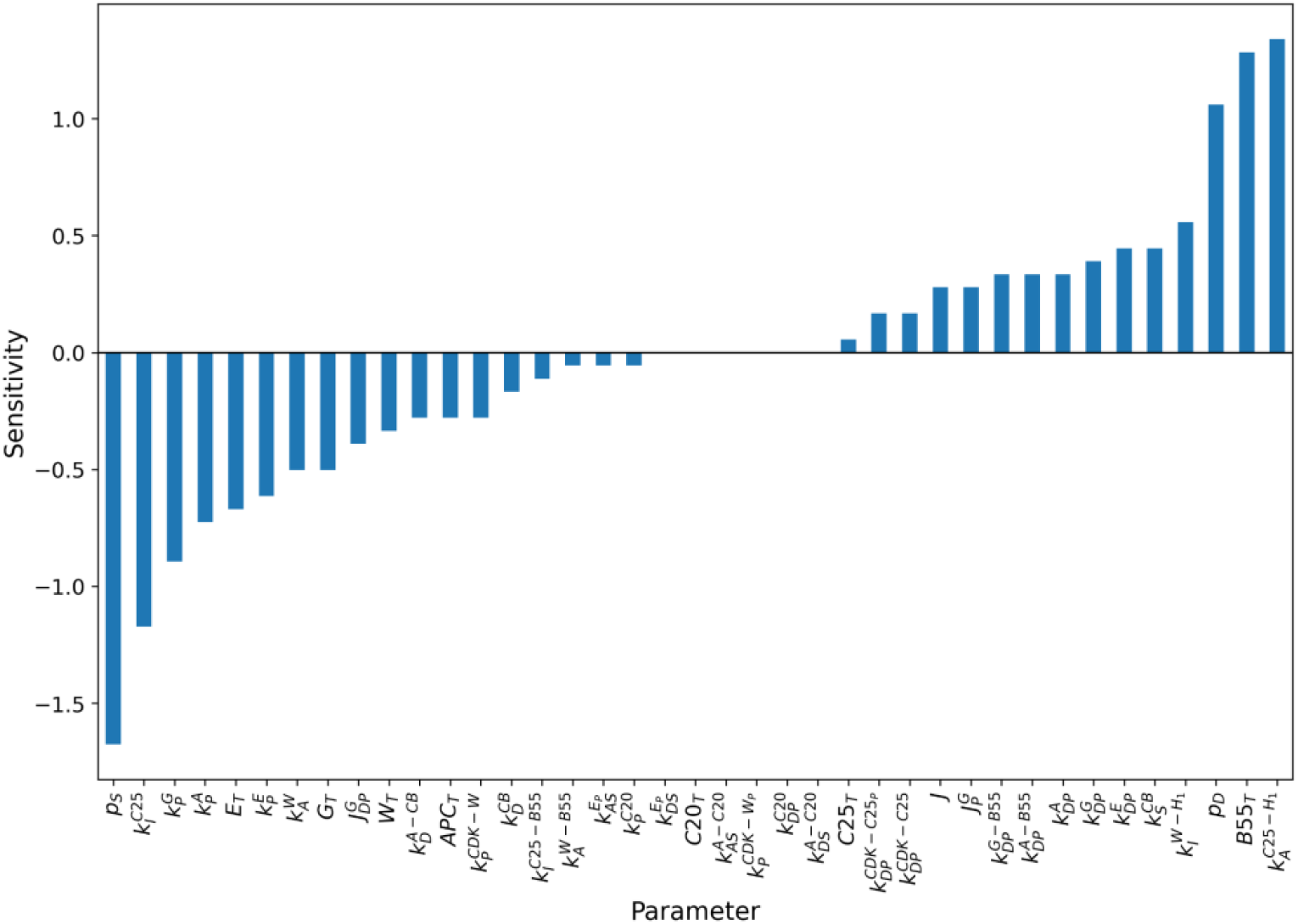
Parameter sensitivity analysis. Each parameter was increased by 1% and the percentage change in dilution threshold was calculated as the sensitivity. Positive sensitivity values indicate an increase in robustness while negative values indicate a decrease.

**Snapshot of Movie S1:**
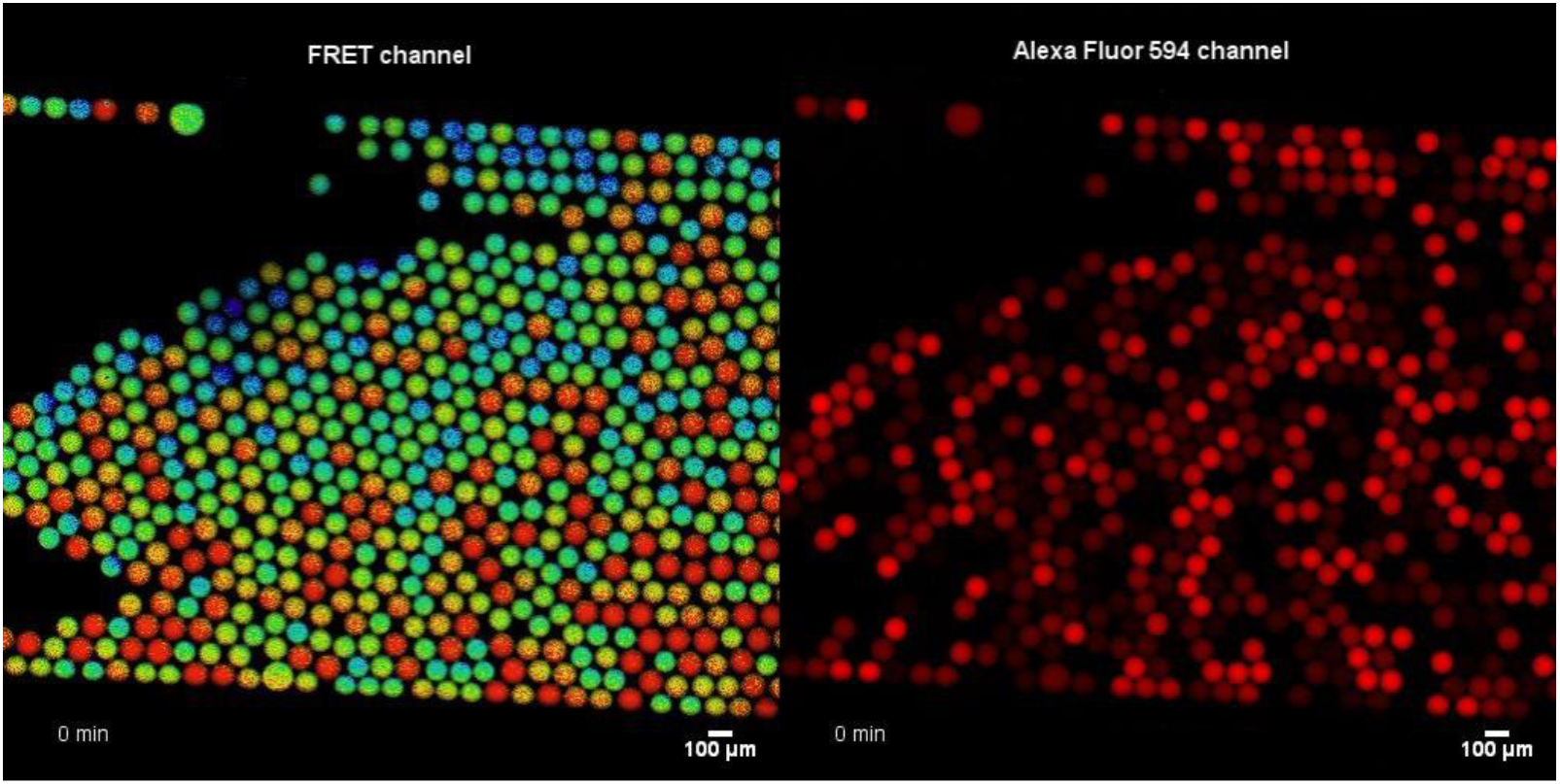
This video is corresponding to Fig 1B. The oscillation behaviors of droplets with different dilutions are recorded. CDK1-FRET sensor is added to indicate the active CDK1 level in the extracts encapsulated in each droplet. In the FRET channel, droplets in red indicated high active CDK1 levels as in mitosis, whereas cool blue color was associated with low active CDK1 levels as in interphase. Droplets with intact mitotic oscillators were supposed to alternate from high to low CDK1 states, resulting in droplets blinking over time. Alexa fluor 594 dextran dye is added as a dilution percentage indicator. Scar bar is 100 μm.

**Snapshot of Movie S2:**
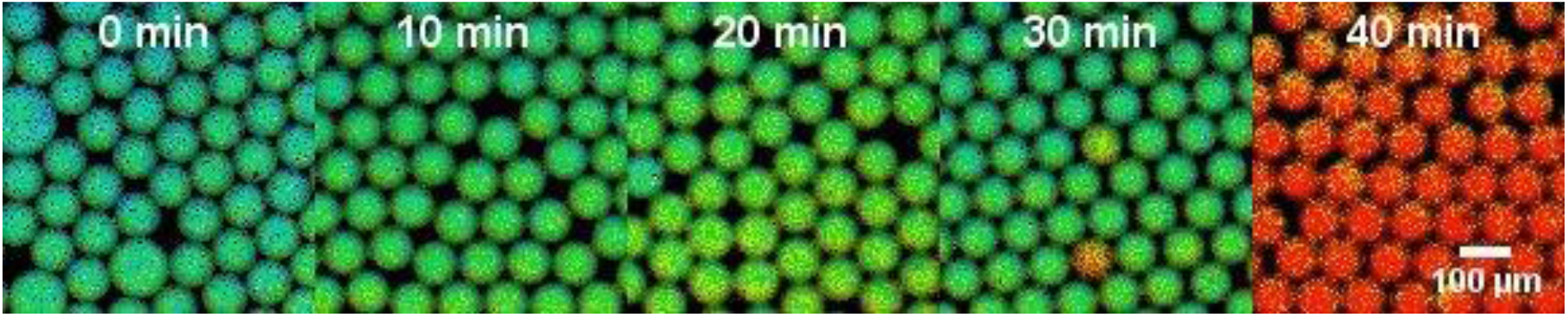
This is corresponding to the concentration experiment in Fig 2A-B. FRET channels for extracts concentrated with different evaporation time are presented. Extracts that are concentrated by vacuum evaporation for 0, 10, 20, 30 minutes still present intact oscillation behaviors, but the 40-min concentrated system stops oscillating.

**Snapshot of Movie S3:**
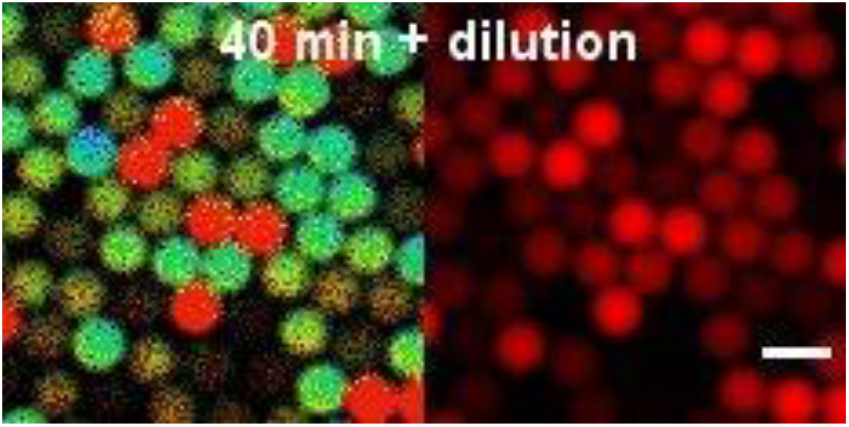
This is corresponding to the diluting of concentrated extract experiment in Fig 2B-I (Left: FRET channel for cell cycle monitoring; Right: Alexa fluor 594 channel for dilution percentage indication). Dilution of non-oscillatory 40-min concentrated extract restores oscillations. Scale bar is 100 μm.

## SI text S1: Mathematical model and dependency of parameters with cytoplasmic density

We began our analysis using as reference a previously developed model of cell cycle dynamics (1). This model describes the regulation of Cdk1:CyclinB complex by Cdc25 and Wee1, the regulation of PP2A by GWL and ENSA, and the negative feedback loop composed of Cdc20 and APC/C. A diagram of the interactions between components is presented in the main text Fig 3A. To model the effect of a variable cytoplasmic density on the system’s parameters we introduce a new parameter *d*, the relative cytoplasmic density. A value *d* = 1 of corresponds to the unperturbed cell cycle dynamics described by the equations of (1). The equations describing the dynamics of the system at different cytoplasmic densities are the following (newly introduced parameter d is shown in red for visual clarity):

**- Cyclin B**

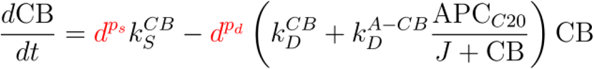

**- Cdk1:CycB complex**

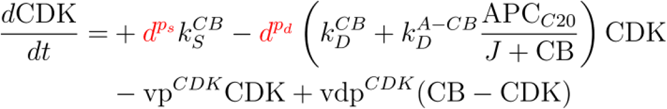

**- Cdc25**

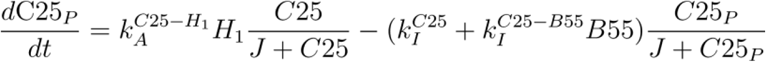

**- Wee1**

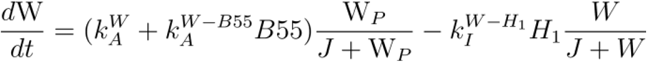

**- PP2A:B55 complex**

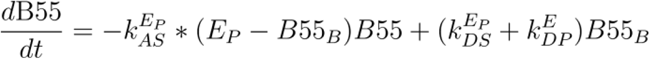

**- Ensa**

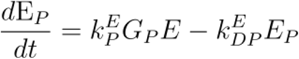

**- Greatwall**

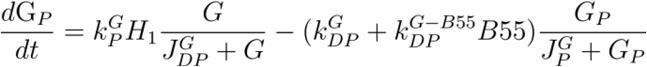

**- Phosphorylation steps for APC partial activation**

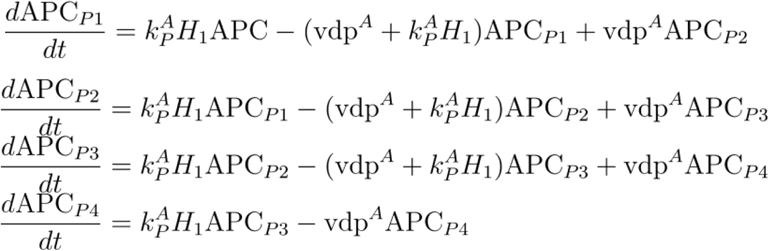

**- APC:Cdc20 complex**

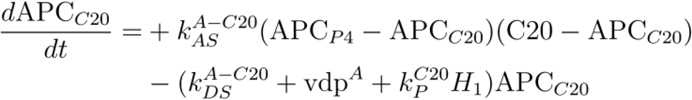

**- Cdc20**

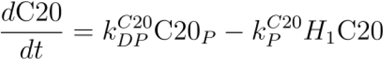

**Auxiliary quantities**

Cdk effective concentration for kinase activity

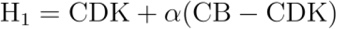

Unphosphorylated Cdc25

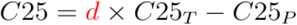

Phosphorylated Wee1

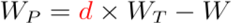

CDK’s phosphorylation rate

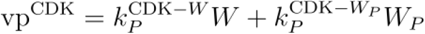

CDK’s dephosphorylation rate

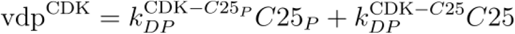

B55 bound to Ensa

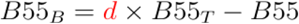

Unphosphorylated Ensa

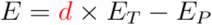

Unphosphorylated Greatwall

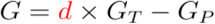

Unphosphorylated APC

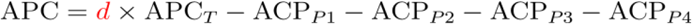

APC’s dephosphorylation rate

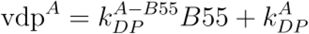

Phosphorylated Cdc20

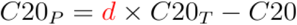

Where we explicitly model the effect of cytoplasmic density on the system’s parameters. First, we assumed that reaction rates of association/dissociation and phosphorylation/dephosphorylation do not depend on cytoplasmic density. Our reasoning is that changes in these microscopic rates will be less important than the changes due to variations in the total concentration of reactants. We realize that changes in viscosity and the properties of the medium can affect these microscopic rates but we chose not to consider those in our first approximation to the phenomenon. Similarly, we assume that Michaelis constants do not vary with cytoplasmic density since in turn these are described by microscopic rates. Second, we assumed that total concentrations scale linearly with cytoplasmic density because experimentally only buffer is added to/removed from the system and not cell cycle’s components. Finally, we considered the effect of changes in cytoplasmic density on synthesis and degradation rate of cyclin B. These rates are expected to be affected by dilution and concentration of the system. Each reaction is composed of multiple intermediary steps which are not explicitly described in the proposed model. Both processes are described effectively with synthesis having a constant production rate and degradation as a first-order reaction. We maintained these simplified descriptions of both processes and explored different hypotheses for possible dependencies with cytoplasmic density. Our guide was the observed changes in cell cycle dynamics due to dilution and concentration. We assumed that synthesis and degradation are scaled by a power of the relative cytoplasmic density denoted p_s_ and p_d_ respectively. Both are undetermined parameters and in SI text S2 we discuss our methodology for the selection of their values.

### SI text S2: Parameter values

The parameters used in our article for the description of physiological cycles slightly differ from those in (1). Specifically, we employed a different synthesis rate since bifurcation diagrams with this higher value of the parameter agreed better with experimental observations. Additionally, the timescale of the system was redefined to match the average observed physiological period in experiments. The timescale was redefined by scaling all reaction rates by a factor of 0.5. The reference physiological parameters used in this study are presented in Table S2.

**Table S2:**
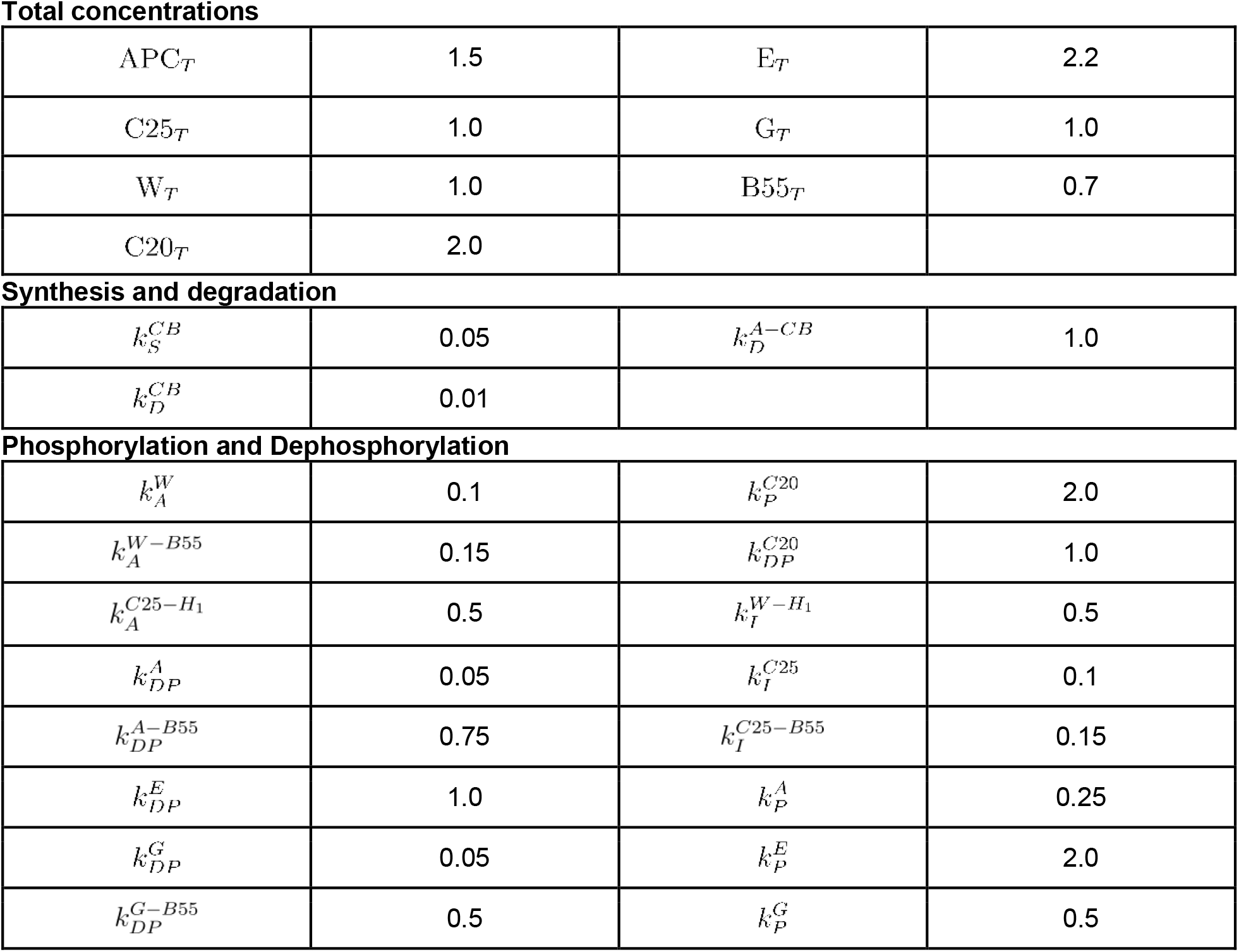
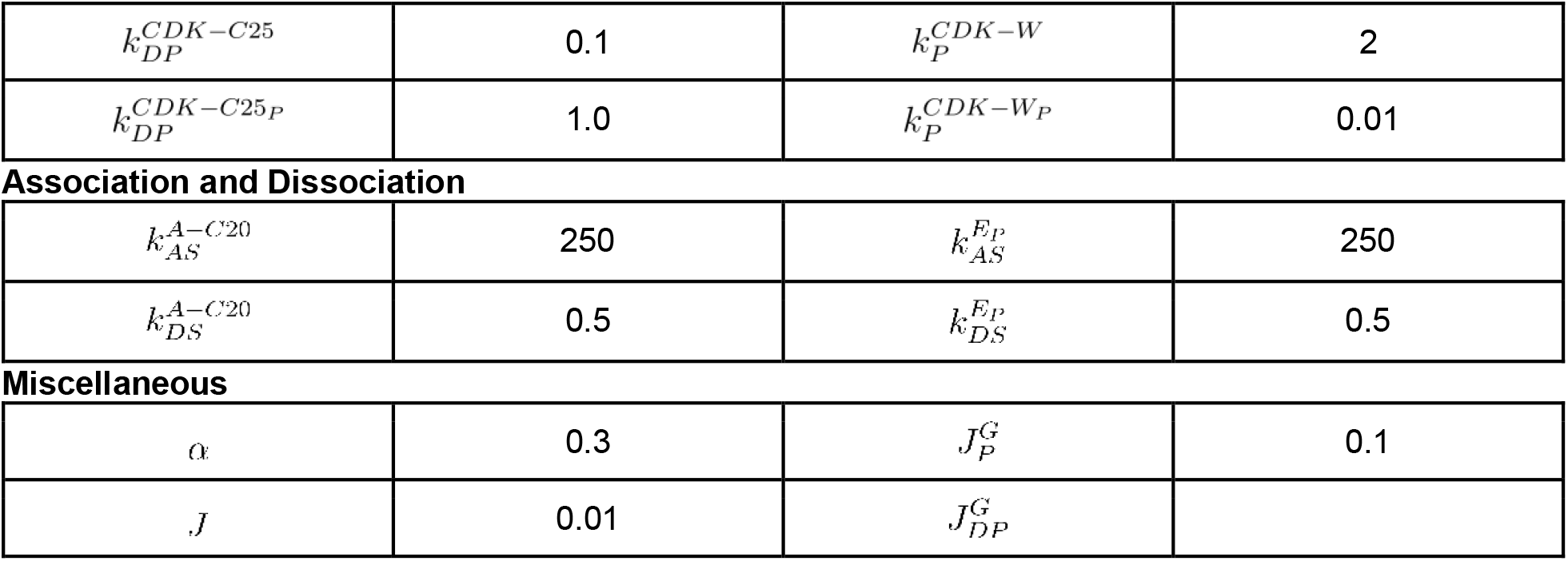
Parameters of the mathematical model describing cell cycle oscillations.

Total concentrations
|  |  |  |  |
| --- | --- | --- | --- |
| $APC_T$ | 1.5 | $E_T$ | 2.2 |
| $C25_T$ | 1.0 | $G_T$ | 1.0 |
| $W_T$ | 1.0 | $B55_T$ | 0.7 |
| $C20_T$ | 2.0 | | |
| Synthesis and degradation |  |  |  |
| $k_S^{CB}$ | 0.05 | $k_D^{A-CB}$ | 1.0 |
| $k_D^{CB}$ | 0.01 | | |

|  |  |  |  |
| --- | --- | --- | --- |
| $k_A^W$ | 0.1 | $k_P^{C20}$ | 2.0 |
| $k_A^{W-B55}$ | 0.15 | $k_{DP}^{C20}$ | 1.0 |
| $k_A^{C25-H_1}$ | 0.5 | $k_I^{W-H_1}$ | 0.5 |
| $k_{DP}^A$ | 0.05 | $k_I^{C25}$ | 0.1 |
| $k_{DP}^{A-B55}$ | 0.75 | $k_I^{C25-B55}$ | 0.15 |
| $k_{DP}^E$ | 1.0 | $k_P^A$ | 0.25 |
| $k_{DP}^G$ | 0.05 | $k_P^E$ | 2.0 |
| $k_{DP}^{G-B55}$ | 0.5 | $k_P^G$ | 0.5 |
| $k_{DP}^{CDK-C25}$ | 0.1 | $k_P^{CDK-W}$ | 2 |
| $k_{DP}^{CDK-C25P}$ | 1.0 | $k_P^{CDK-WP}$ | 0.01 |
| **Association and Dissociation** |  |  |  |
| $k_{AS}^{A-C20}$ | 250 | $k_{AS}^{EP}$ | 250 |
| $k_{DS}^{A-C20}$ | 0.5 | $k_{DS}^{EP}$ | 0.5 |
| **Miscellaneous** |  |  |  |
| $\alpha$ | 0.3 | $J_P^G$ | 0.1 |
| $J$ | 0.01 | $J_{DP}^G$ | |

Where we keep the same convention as in (1) of concentrations being in arbitrary units and time being represented in minutes. We have two undetermined parameters in our model, *p_s_* and *p_D_*. They represent the changes in synthesis and degradation rate as the cytoplasmic density is modified. As an example, if protein synthesis was simply described as a second order reaction between total mRNA and ribosomes, we would expect *ps* = 2 to account for the decay of both total concentrations. Similarly, if degradation was effectively a reaction between the proteasome and the substrate, we would expect *p_D_* = 1 to account for the scaling of proteasome concentration. However, given that both synthesis and degradation are mediated by multiple intermediaries, we decided to explore different values of p_s_ and p_d_. We considered all possible integer combinations between 0 and 4 for both parameters. For each combination we compared experiments and simulations in terms of the period of the oscillations, the amount of time spent in the rising phase and the amplitude. Using these three features we filtered out those parameter combinations that did not follow experimental trends.

We considered specific experimental observations to select the most appropriate model. First, we expect the period to increase as a function of dilution and oscillations to stop in a range between 60% to 80%. Second, close to the threshold rising periods become elongated, which translates to an increase in the fraction of rising period to total period. Third, amplitude should decay with dilution and display a finite value at the threshold in contrast to decaying continuously to zero. We simulated and measured these quantities for all studied combinations of p_s_ and p_d_. The results can be appreciated in Fig S4. Simulation details are described in SI text S6.

We observed that simulation results could be clustered into three groups. While most simulations show a period that increases with dilution, the behavior for the rising fraction and amplitude is different between simulations with p_s_ = p_d_ + 1, p_s_ > p_d_ + 1, and p_s_ ≤ p_d_ + 1. The only group that meets the three conditions observed in the data is the one where p_s_ = p_d_ + 1. Namely, period increases with dilution, thresholds are between 60% and 80%, rising fraction increases just before oscillations stop, and amplitude decreases without dropping to 0 near the threshold. In contrast the other groups show features not observed in data. The group p_s_ > p_d_ + 1 presents an amplitude that decays continuously to zero with sudden jumps in period near the threshold. The group p_s_ ≤ p_d_ + 1 displays decreasing rising fraction as a function of dilution in disagreement with experimental observations. Therefore, we selected the models with p_s_ = p_d_ + 1 for further analysis We compared the selected simulation curves (p_s_ = p_d_ + 1) with different experimental replicates (Fig S5). A great variability in the response to dilution is observed in experiments regarding the exact period of the oscillations. However, all curves displayed the same trend: increase in period with a sudden stop of oscillations consistent with our selected models. We arbitrarily selected the case p_s_ = 3, p_d_ = 2 to present in the main text but other specific values of p_s_ and p_d_ that satisfy p_s_ = p_d_ + 1 agree with specific experimental days equally well.

### SI text S3: Encapsulation error and droplet simulations

Next, we explored the possibility of droplet-to-droplet variability due to encapsulation effects. It has previously been shown that encapsulation can modify the behavior of oscillators (2). Here we will adopt the same mathematical framework to describe this effect by using a gamma distribution to draw certain parameters in our model and generate a population of *in silico* droplets. We consider total concentrations as well as synthesis and degradation rate to be affected by this unequal partitioning. On the other hand, we do not consider microscopic reaction rates to be affected by this partition. Here, we retain the same working hypothesis that the major changes in reaction rates come from variations in the total concentration of reactants.

The probability density we adopt for sampling our parameters takes the form

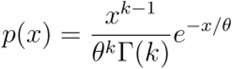

Where x is the parameter being sampled, k the shape of the distribution, θ its scale, and Γ the gamma function. The gamma distribution allows us to set the mean and variance independently. Given that the mean value of x under this distribution is given by kθ we assign

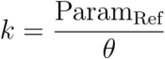

To center the distribution around the corresponding bulk reference parameter. Then, θ becomes a free parameter that describes the variability of the parameter being sampled. From experiments we can estimate the coefficient of variation (std/mean) when encapsulating a fluorescent probe. Estimates taken from experiments in Table S1, produce a CV in the order of 1e-2. Since the standard deviation of the selected distribution is kθ^2^, with our choice of k we have

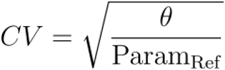

Since model parameters are in relative units, we plug in a value of Param_Ref_ = 1 for estimating θ. With a CV of 1e-2, we estimate θ to be

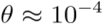

Therefore, to simulate individual droplets, we draw total concentrations, synthesis rate, and degradation rates from a gamma distribution centered on our selected reference parameters (Table S2). For completion, we simulated the system under different values of θ (Fig S6). When compared with the variability and characteristics of the experimental data, a value of θ = 1e-4 seems appropriate to describe the variability between droplets: larger values present a variability not seen in experiments and smaller values follow too closely the bulk solution.

### SI text S4: Wee1 and Cdc25 Inhibitors

To simulate the effect of inhibitors, we introduce additional parameters in the system. The reaction rate of Wee1 and Cdc25 activity on Cdk1 is multiplied by new variables I_W_ and I_C_ respectively. The modified equation for Cdk1 reads:

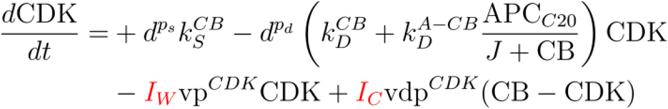

Where a value of I_W_ and I_C_ of 1 represents the system without inhibitors added. By simulating the system for values between 0 and 1 of the new parameters we can mimic the effect of inhibitors that reduce the activity of Wee1 and Cdc25 on Cdk1.

To connect simulation results with experiments we assume a Hill effect of the inhibitors on the enzyme’s activity. Therefore, the inhibition parameters are connected to the concentration of inhibitor via

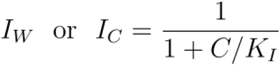

Where C represents the inhibitor concentration and K_I_ the unknown inhibition constant. For Wee1 inhibition and Cdc25 inhibition we simulate a population of droplets for both dilutions (d) and inhibitions (I_W_ or I_C_) between 0 and 1. We process our results to obtain the oscillation percentage, period, and thresholds at each inhibition percentage (Fig S9). Results are converted to inhibitor concentration by exploring different values of K_I_ and selecting the value that resembles experimental observations more closely. For Wee1 simulations we used a value of K_I_ = 0.175 uM. For Cdc25 simulations we used K_I_ = 10 uM.

### SI text S5: Sensitivity analysis

The mathematical model employed in this manuscript has several parameters describing the processes driving cell cycle oscillations. Here, we explore which parameters when changed affect the most the robustness to dilution. We increased each parameter of the model by 1% and measured the dilution at which oscillations disappeared (Threshold 1%). Then we compared that dilution to the reference value when the parameter was not changed (Threshold Ref). This allowed us to calculate the sensitivity as

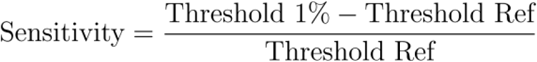

The result is presented in Fig S10. This analysis suggests the most interesting parameters to vary in future experiments. The decay of both synthesis and degradation (p_s_ and p_D_) appear as great contributing factors. Similarly the activation of Cdc25 and Greatwall by Cdk1 appears as important variables when determining the robustness to dilution. Finally, the total concentration of PP2A could be explored to change the system’s robustness. As confirmed by experiments and simulations the activity of Wee1 and Cdc25 towards Cdk1 is not a parameter of the model that when changed greatly affects the robustness to dilution.

### SI text S6: Simulation details and analysis

All time series are obtained by numerical integration of the ODE equations. Simulations were performed in Python 3.7.10 using the function solve_ivp from the package scipy 1.4.1. The selected integration method was LSODA. Oscillatory or steady-state behavior is determined by the norm of the system’s velocity-vector in conjunction with an analysis of the signal’s peak detection features. Namely, the period stability (variability in time differences between detected peaks) and amplitude stability (variability in peak concentrations) are analyzed. Peaks were detected with the function find_peaks from the same scipy package version. Droplet simulations are ODE simulations where the corresponding total concentrations, synthesis rate, and degradation rate are sampled from a Gamma distribution representing the encapsulation error. In all cases, 500 droplets are simulated per relative cytoplasmic density value. For various utilities numpy 1.19.4, pandas 1.2.1, and matplotlib 3.3.3 are used. Bifurcation diagram in 3I is performed in XPPAUT 8.0. The diagram is started from a high dilution converged steady state. Stable and unstable steady states are first tracked while the Hopf bifurcations are continued in a second iteration of the algorithm. AUTO parameters used for the scanning are: Ntst = 60, Ds = 0.01, Dsmin = 0.001, Dsmax = 0.1, ParMin = 0, ParMax = 2.5. Remaining parameters are set to their default values. In all cases except for the blue curve in 3J, all simulations started from the same initial condition as in (1) rescaled by the appropriate dilution. The blue curve in 3J uses as the initial condition the steady state of the system when the relative cytoplasmic density is 2.5.

